# Common neural and transcriptional correlates of inhibitory control underlie emotion regulation and memory control

**DOI:** 10.1101/708552

**Authors:** Wei Liu, Nancy Peeters, Guillén Fernández, Nils Kohn

## Abstract

Inhibitory control is crucial for regulating emotions, and it may also enable memory control. However, evidence for their shared neurobiological correlates is limited. Here, we report meta-analyses of neuroimaging studies on emotion regulation, or memory control, and link neural commonalities to transcriptional commonalities using the Allen Human Brain Atlas (AHBA). Based on 95 fMRI studies, we reveal a role of the right inferior parietal lobule embedded in a frontal-parietal-insular network during emotion and memory control, which is similarly recruited during response inhibition. These co-activation patterns also overlap with the networks associated with “inhibition”, “cognitive control”, and “working memory” when consulting the Neurosynth. Using the AHBA, we demonstrate that emotion and memory control-related brain activity patterns are associated with transcriptional profiles of a specific set of “inhibition-related” genes. Gene ontology enrichment analysis of these “inhibition-related” genes reveal associations with the neuronal transmission and risk for major psychiatric disorders as well as seizures and alcoholic dependence. In summary, this study identified a neural network and a set of genes associated with inhibitory control across emotion regulation, memory control. These findings facilitate our understanding of the neurobiological correlates of inhibitory control and may contribute to the development of novel brain stimulation and pharmacological interventions.

## Introduction

One of the most influential cognitive theories of emotion regulation proposes that it is primarily supported by inhibitory control (Ochsner and Gross, 2005; Gross and Thompson, 2007). A potentially related cognitive process, memory control, is defined as an ability to actively reduce the accessibility of memories. Memory control is also thought to be supported by inhibitory control (Anderson and Green, 2001; Anderson *et al.*, 2004; Anderson and Hanslmayr, 2014). Recently, Engen and Anderson summarized the conceptual link between emotion regulation and memory control and raised the question of whether there is a common neurobiological mechanism supporting these two processes (Engen and Anderson, 2018). Here, we set out to empirically demonstrate the neurobiological commonalities between the two processes by first analyzing fMRI data and then linking neuroimaging findings with post mortem gene expression data.

The idea that inhibitory control, as the fundamental cognitive process, supports other higher-level processes such as emotion regulation and memory control is supported by behavioral and neural evidence. Behaviorally, a positive correlation between emotion regulation and memory control performances, has been found (Depue *et al.*, 2016). Human neuroimaging results showed overlapping recruitment of right superior medial frontal gyrus (rSFG), right inferior frontal gyrus (rIFG) in both emotion regulation (Ochsner *et al.*, 2002; Buhle *et al.*, 2014; Kohn *et al.*, 2014) and memory control (Anderson *et al.*, 2004; Guo *et al.*, 2018). Beyond these frontal regions, the parietal cortex is another candidate region that may play a critical role in both emotion regulation and memory control. Starting from the idea of multiple-demand system in the brain (Duncan, 2010), both the evidence from task-fMRI (Fedorenko *et al.*, 2013) and resting-state fMRI (Dosenbach *et al.*, 2007; Power *et al.*, 2011) suggest that a fronto-parietal network supports the initiation of top-down control and control adjustment in response to task-goals and feedbacks.

No previous formal meta-analysis of human neuroimaging studies has investigated the neural commonalities between emotion regulation and memory control to pinpoint the overlap in inhibitory control, although common neural correlates between emotion regulation or memory control with response inhibition tasks (e.g., stop-signal and Go/Nogo task) have been investigated in two recent meta-analyses (Guo *et al.*, 2018; Langner *et al.*, 2018). Using Activation likelihood estimation (ALE), we aim to demonstrate that emotion regulation and memory control evoke activation of similar brain regions with a spatial pattern that is similar to the activation of typical response inhibition paradigms, including stop-signal and go/no-go tasks. Moreover, beyond overlapping regional brain activity, we also are interested in overlapping co-activation patterns of associated brain regions. We used meta-analytic connectivity modeling (MACM) to test whether these brain regions act as an interconnected functional network. We expected to find a similar set of co-activated brain regions that are associated with both emotion regulation and memory control, defining a core “inhibition-related” network.

Identifying an underlying neural network is relevant for potentially brain stimulation interventions including but not limited to Transcranial Magnetic Stimulation (TMS) and Transcranial Direct Current Stimulation (tDCS). However, further dissection of molecular underpinnings is necessary to expand our understanding of inhibition control, which in turn may pave the way for pharmacological interventions. Therefore, we reasoned whether emotion regulation and memory control are not only related to a common neural network defined by activity and connectivity, but also by associated similarities in spatial transcriptional profiles. Conventional imaging-genetic methods including candidate gene methods and genome-wide association approaches cannot reveal the relationship between localized gene transcription and task-related brain activity, even though they already showed the association between multiple common gene variants and neuroimaging measures (Hariri *et al.*, 2002; Elliott *et al.*, 2018). Thus, to explore the relationship between spatial transcriptional profiles and emotion regulation or memory control-related brain activity, we adopted a recently developed approach to associate spatial maps of gene expression in post-mortem brain tissues with brain activation measures (Gorgolewski *et al.*, 2014; Kong *et al.*, 2016). Spatial pattern analysis of gene expression maps of the Allen Human Brain Atlas (AHBA) together with neuroimaging revealed fundamental features of transcriptional regulation (See review by Fornito, Arnatkevičiūtė, & Fulcher, 2018) and related disruptions in brain disorders (Romme *et al.*, 2017; Grothe *et al.*, 2018; McColgan *et al.*, 2018; Romero-Garcia *et al.*, 2018). However, the correspondence between spatial transcriptional profiles and neural functionality has not yet provided full insight into how genetic correlates are linked to core cognitive abilities (e.g., inhibitory control in this study). Based on the assumption that spatial transcriptional profiles not only co-vary with the connectional architecture but also support the task-evoked, synchronous brain activity (Gorgolewski *et al.*, 2014; Kong *et al.*, 2016; Berto *et al.*, 2017; Shine *et al.*, 2019) we expected to find similar spatial transcriptional profiles between emotion regulation, memory control, and response inhibition.

To investigate the neural and transcriptional commonality of emotion regulation and memory control, we combined task fMRI data, neuroimaging meta-analytic approaches, and postmortem gene-expression data. First, we used the activation likelihood estimation method to generate the task activation map for each paradigm of interest (e.g., emotion regulation or memory control) based on 95 published task fMRI studies with in total 1995 healthy participants (Figure 1A). Then, we used the brain regions identified by the ALE as seed regions and estimated the meta-analytic connectivity map (or co-activation map), using information from BrainMap (Figure 1B). And to explore the behavioral relevance of these findings, we examined the associations of these co-activation patterns in a data-driven way via Neurosynth with behavioral domains (Figure 1C). Next, we calculated the spatial association between task activation maps and human gene expression maps to identify their common spatial transcriptional profiles (Figure 1D). Finally, we performed a systematic and integrative analysis of the resulting gene list to gain an in-depth understanding of putative biological functions and disease associations of the identified “inhibition-related “gene set (Figure 1E).

**Figure 1.**
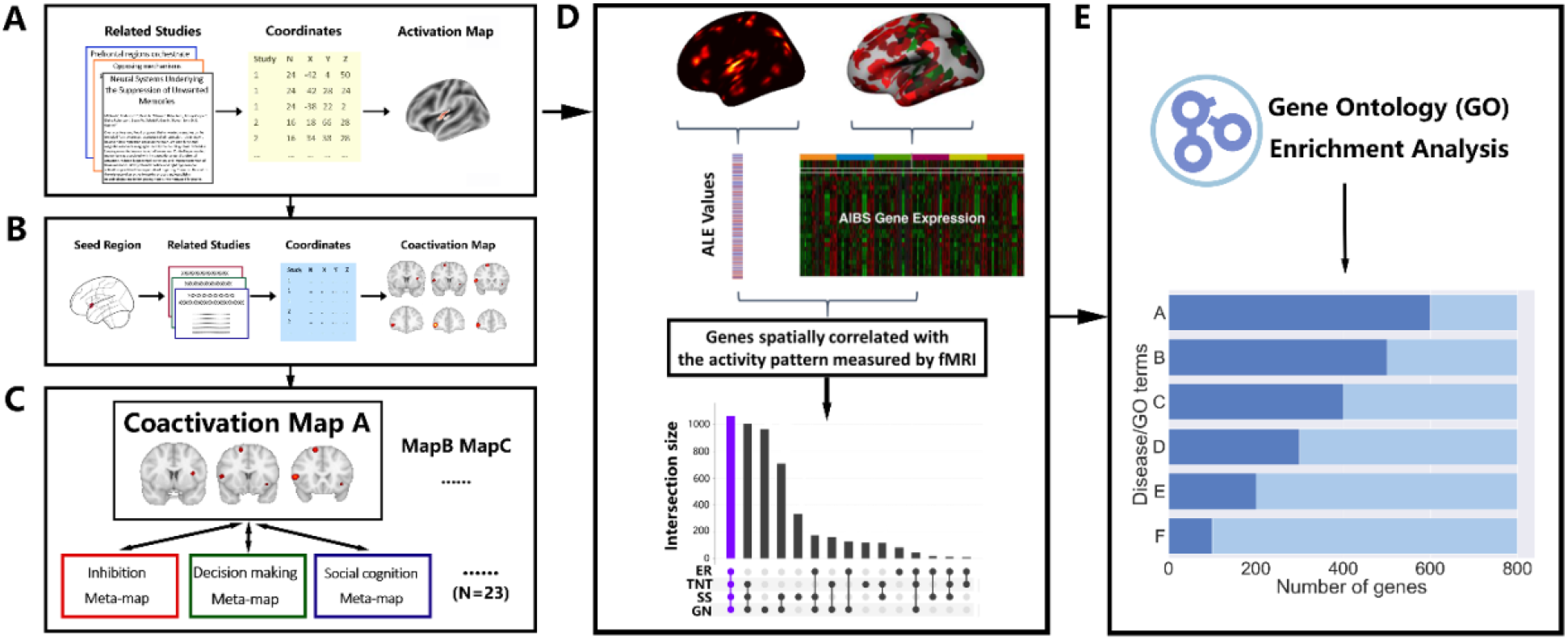
Schematic of the pipeline for the investigation of neural and transcriptional commonality. (A) Activation likelihood estimation (ALE) meta-analyses of functional MRI studies. MRI coordinates of reported brain activations from relevant studies were used to generate the activation map for each task of interest. (B) Meta-analytic connectivity modeling of seed regions resulting from the ALE analyses was conducted to search co-activated regions over all studies in BrainMap. Coactivation maps were estimated via ALE. (C) To generate behavioral profiles of co-activation maps, we used Neurosynth to acquire a list of meta-maps, including the behavioral terms from low-level to high-level cognitive processes (total Number of terms=23). Next, each co-activation map was compared to all meta-maps (e.g., inhibition, decision making, and social cognition) to compute the similarity index for each meta-map. (D) The activation patterns were associated with gene expression maps from the Allen Human Brain Atlas (AHBA). Each gene expression map was vectorized based on the expression measures within the three-dimension maps. The ALE values at the corresponding brain regions on the activation map were extracted and vectorized. To identify the genes whose spatial patterns are similar to a specific activation map, the similarity between all vector pairs were quantified. “Inhibitory-related” genes were defined by identifying genes of which the expression patterns are correlated with the activation maps of four paradigms (ER, Emotion Regulation; TNT, Think/No-Think; SS, Stop-signal; GN, Go/No-go). (E) “Inhibition-related” genes were associated with biological functions (GO terms) and disease terms for the interpretation using Gene Ontology Enrichment Analysis (GOEA).

## Materials and Methods

### Literature searches, selection, and coordinates extraction

In total, we perform literature searches for four task paradigms (think/no-think, emotion regulation, stop-signal, and go/no-go) and one network (“default mode network”). To avoid biases, we use the same inclusion criteria during the search.

1. We only included data from studies on healthy adults with no prior report of neurological, medical, or psychiatric disorders in the current meta-analysis, while results of patients or specific sub-group effects (e.g., gender differences) were not included. Articles including patients were only selected if they reported results for a control group separately, and only the control group was included here.
2. Only neuroimaging studies, which used whole-brain fMRI and reported coordinates for brain activation or deactivation in standard anatomical reference space (Talairach/Tournoux; Montreal Neurological Institute (MNI)) were considered. Coordinates originally published in Talairach space were converted to MNI space using the algorithm implemented in GingerALE 3.0.2 (http://www.brainmap.org/ale/)
3. Only studies reporting results of General Linear Model (GLM) contrasts were included, while studies focusing on functional connectivity, structural, resting-state, or brain-behavior correlations were excluded.
4. Only studies reporting whole-brain analyses were included, while studies based on partial coverage or employing only region-of-interest analyses were excluded.

Detailed search and extraction procedures were as following for each paradigm:

#### Think/No-Think studies

A step-wise procedure was used to search articles, published before February 2020, using functional MRI to investigate brain activity during Think/No-Think paradigm. First, we used standard search in PubMed and ISI Web of Science to perform the search. More specifically, we used the combination of following key words during the search: “memory regulation”, “memory control”, “memory suppression regulation”, “memory inhibition”, “think/no-think”, “fMRI”, “neuroimaging”, “functional magnetic resonance imaging”, or “functional MRI”. At the same time, we carefully exclude studies using the “directed forgetting” paradigm, which targets at the memory control during the encoding. Next, two lab members compared the search results with a recent review article (Anderson and Hanslmayr, 2014) to find additional relevant studies. The same two lab members independently extracted the coordinates and other essential information (e.g., sample size, type of stimulus) extraction based on the identified think/no-think literature and then cross-validated the coordinates. In summary, this search and inclusion/ exclusion criteria led to 15 think/no-think studies (491 subjects and 256 foci).

##### Emotional Regulation Studies

We take the databases of previously published meta-analyses on emotion regulation (Kohn *et al.*, 2014; Morawetz *et al.*, 2017). We use the keywords:“emotion regulation”, “affective regulation”, “implicit emotion regulation”, “explicit emotion regulation”, “interpersonal emotion regulation”, “extrinsic emotion regulation, “intrinsic emotion regulation”, “reappraisal”, “suppression”, “distraction”, “detachment”, “labelling”, “affective labelling”, “reinterpretation”, “rumination”, “fMRI”, “neuroimaging”, “functional magnetic resonance imaging”, or “functional MRI”. In the case that a study did not report the contrast of interest for this meta-analysis, the corresponding authors were contacted and asked to provide more information on their data. In the following the term “experiment” refers to any single contrast analysis, while the term “study” refers to a scientific publication, usually reporting several contrasts, i.e. experiments. This search and the employed inclusion/exclusion criteria led to a total inclusion of 107 studies from peer-reviewed journals by July 31st, 2017 (385 experiments, 3204 participants).

Each experiment was manually coded by the authors of the previous meta-analysis (C.M. and N.K) with terms that described the experimental design with respect to contrast, stimulus type utilized, emotion regulation strategy, goal of the strategy, valence of the stimuli, tactics of the strategy and the task nature. To achieve a more appropriate comparison between think/no-think and emotion regulation, we restrict inclusion to ER studies that used the “suppression” or “distraction” strategy. This led to a similar amount of studies compared to memory control and most crucially suppression or distraction of emotions is conceptually also closer to the process of suppressing memories. These criteria led to the inclusion of 15 emotion regulation studies (387 subjects and 165 foci).

##### Go/No-go and Stop-signal studies

A similar procedure was used to search published whole-brain functional MRI studies using the “Go/No-go” and “stop-signal” paradigm. To confirm the completeness of our search, we compared our results with used studies in a recent meta-analysis of motor inhibitory and memory control (Guo *et al.*, 2018). Again, two lab members (W.L and N.P) performed the coordinates and study information extraction for the Go/No-go and Stop-signal studies.

##### Default Mode Network (DMN) (task-negative network) identification

We also performed a coordinated-based meta-analysis to identify the DMN. Instead of manually searching related studies and extracting coordinates, we used the BrainMap database (Fox and Lancaster, 2002; Laird *et al.*, 2005) to find peak coordinates of task-independent deactivation reported in neuroimaging literature. This method was used before by Laird and colleagues (A. R. Laird *et al.*, 2009) to identify the core regions in DMN. More specifically, we searched the BrainMap for all contrasts that were labeled as “deactivation” and “low-level control” during submission. “Deactivation” refers to contrasts in which stronger signal was observed during a baseline condition than during task condition (e.g., Control-Task); “low-level control” are conditions in which either fixation or resting was defined as the baseline. Our search was further limited to “normal mapping”, which means that participants who diagnosed with disease or disorders were excluded. In total, 105 studies (1588 foci) matched our search criteria and were used in the following analyses.

### Activation Likelihood Estimation (ALE) analyses

The ALE analyses were based on the revised ALE algorithm (Eickhoff *et al.*, 2009) in GingerALE 2.3. Firstly, two separate meta-analyses were conducted for the Think/No-Think (contrast: No-Think vs. Think) and emotion regulation (contrast: Regulation vs. View) tasks using cluster-level inference (FWE cluster-level correction p<0.05, uncorrected cluster-forming threshold p<0.001, threshold permutations=1000). Secondly, contrast analyses (Eickhoff *et al.*, 2011) were conducted between the Think/No-Think and emotion regulation tasks. In these contrast analyses, the thresholded activation maps from the two separate analyses, as well as the pooled results from both tasks, were used as inputs. Conjunction and contrast maps between the conditions were given as output. For the output images, the same cluster-level threshold correction was used (FWE cluster-level correction p<0.05, uncorrected cluster-forming threshold p<0.001, threshold permutations=1000).

Additionally, meta-analyses of published fMRI studies using the Stop-signal and Go/No-go paradigms were also performed. The same software (GingerALE 2.3) and threshold (cluster-level p<0.05) was used to perform the analysis. Due to the unbalanced number of studies and subjects included, no conjunction or contrast analyses were performed for four tasks to identify the overlap.

### Co-activation analyses using BrainMap

We conducted the Meta-Analytic Connectivity Modeling (MACM) analyses on the regions from the ALE meta-analysis. More specifically, for each ROI, we used the BrainMap database (Angela R Laird *et al.*, 2009; Laird *et al.*, 2011) to search for experiments that also activated the particular ROI. Next, we retrieved all foci reported in the identified experiments. Finally, ALE analyses were performed over these foci to identify regions of significant convergence. Sequentially, raw co-activation maps were corrected for multiple comparisons (Voxel-wise False Discovery Rate(FDR)<0.05. All clusters size> 200mm^3^)

### Functional profiles of the co-activation maps

To assess associated functional terms of the co-activation maps generated by MACM, we used the NeuroSynth meta-analytic database (www.neurosynth.org) (Yarkoni *et al.*, 2011). We followed the methodology (See Meta-analytic Functional Gradients section) used in a previous study to assess topic terms associated with the principal connectivity gradient in the human brain (Margulies *et al.*, 2016). More specifically, we conducted a term-based meta-analysis for the same list of NeuroSynth topic terms as Margulies and colleagues did. This list covered well-studied functional terms from low-level cognition (e.g., visual perception, and auditory) to high-level cognition (e.g., language, and rewarding). Sequentially, we examined the association between these term-based activation maps with our four sets of co-activation maps (emotion regulation, think/no-think, stop-signal, go/no-go). For each co-activation map, a spatial similarity index (r statistic) between each co-activation map and each meta map of the functional term was provided. The terms were then ordered based on the average correlation for interpretation and visualization.

### Activation-gene expression association analysis

We used a recently developed activation-gene expression association analysis to link the task-related brain activity to gene expression in postmortem human brains. This analysis can identify a list of associated genes based on the MRI-space statistical map(s). This analysis presupposes that if certain gene(s) are associated with the cognitive task of interest, then spatial distributions of their expression values and task-related activation pattern measured by functional MRI should be similar.

The Allen Human Brain Atlas (http://www.brainmap.org) was used in the gene expression decoding analysis. The atlas provided genome-wide microarray-based gene expression data based on six postmortem human brains (Gene expression level of over 62000 gene probes for around 1000 sampling sites across the whole brain were recorded) (Hawrylycz *et al.*, 2012). Additionally, structural brain imaging data of each donor was collected and provided, which enable users to visualize gene expression in its naive space and perform the registration to the standard MRI MNI space.

Previous studies used slightly different statistical methods to associate task-independent MRI-based brain measures (e.g., cortical thickness, functional or structural networks) with the gene expression data (Richiardi *et al.*, 2015; Wang *et al.*, 2015; Seidlitz *et al.*, 2018). We used the method developed by Gorgolewski and colleagues implemented in the alleninfo tool (Gorgolewski *et al.*, 2014) (https://github.com/chrisfilo/alleninf). This method was originally designed for the association analysis between voxel-vise statistical maps and gene expression maps. This method was also by default implemented in Neurovault (Gorgolewski *et al.*, 2015) (https://neurovault.org/). The method has two important features: (1) nonlinear coregistration of the donor’s brain with MNI space was allowed and (2) the ability to use a random-effects model makes it possible to generalize the results to the whole population. The activation-gene expression association analysis works as following: (1) data from gene probes were aggregated for each gene, resulting in 20787 gene expression maps. (2) For each gene expression map, MNI coordinates of each sampling location (the locations in which brain tissues were analyzed for the gene expression data) were extracted to draw a spherical ROI (r=4mm). We used these ROIs to extract the average values of the ALE statistical map within each ROI. Next, the gene expression and meta-analysis vectors were correlated. (3) This extraction and correlation procedure was repeated for each gene expression map to quantify the spatial pattern similarity between the statistical maps and gene expression map. (4) Different thresholds (i.e., 500, 1000, 1500, 2000) was implemented to generate the significantly associated gene list(s) among the 20787 genes. We mainly presented results under the threhold of 2000 most significant genes, but results under other thresholds can be found in the Supplemental Material. Since the negative correlation between brain measures and gene expression is difficult to explain, we only considered the positively correlated genes. Additionally, because of fundamental differences in gene expression between cortical and subcortical regions, we only performed our analyses within the cortical regions.

The described association analysis takes MRI statistical map(s) as input(s) and will output a list of significantly associated genes. To investigate the common transcriptional signatures (associated genes) of emotion regulation and memory control, we used the unthresholded statistical maps from the ALE analyses to identify the task-associated gene list via the association algorithm. Next, the common gene list was generated by overlapping the emotion regulation-associated gene list and memory control-associated gene list. To further investigate whether the two gene lists significantly overlapped with each other, **we** generated the null distribution of the number of overlapping genes by creating two gene lists (with the same size as the real gene lists) from the 20787 genes. To control for autocorrelation, we generated 5000 lists in this way and for each we estimated the overall spatial similarity between identified genes (i.e., emotion regulation-related and memory control-related genes) for each donor (Table S3). Identified genes demonstrated modest spatial correlations (mean=0.063, standard deviation=0.27) across different thresholds and donors. Based on these correlation values, 1718 out of 5000 pairs of randomly generated gene lists with a similarity ranging from 0.05 to 0.07, and the standard deviation ranging from 0.25 to 0.29 were further selected. The significance of overlapping genes was estimated by comparing the real number of overlapping genes with the number of overlapping genes within these 1718 pairs.

An alternative method (i.e., “permutated” statistical maps) was also used to evaluate the significance of the overlap further. We permutated the spatial distribution of the fMRI statistical maps used in the activation-gene expression association analysis for 100 times. Voxel-wise statistical values that were stored at the real maps (i.e., emotion regulation or memory control map) were relocated to other locations, testing for spatial specificity of the original task-related spatial patterns, but containing all voxel-wise values. It is notable that this relocation procedure was restricted by the functional-defined brain parcellation (Schaefer *et al.*, 2017). In this way, intrinsic connectivity associations that might underlie functional networks and brain parcels are retained. These “permutated” statistical maps were used for the activation-gene expression association analysis to generate 100 pairs of gene lists. Before the overlapping calculation, we also calculated overall spatial similarity with each pair. Seventy-two out of 100 pairs of these gene lists had a similarity ranging from 0.05 to 0.07 (mean=0.06) with a standard deviation ranging from 0.25 to 0.29 (standard deviation=0.27). These results were used as the null distribution to estimate the p-value of the overlap. To identify the “inhibition-related” genes, we first used the same activation-gene expression association analysis to identify the stop-signal and go/no-go-associated gene list and then defined the overlap between four gene lists (emotion regulation, think/no-think, stop-signal, and go/no-go) as “inhibitory-related” genes.

We aim to improve the specificity of our activation-gene expression association analysis and safeguard the possibility that these identified genes may only support the general functional brain network architecture instead of particular cognitive functions. To rule out this possibility, we performed a dedicated control analysis: (1) An unthreholded DMN ALE map was used as a comparison with four inhibitory control tasks because control processes hardly take place in the DMN and the DMN has been associated with homeostasis and undirected thought or mind-wandering. (2) A pair of “permutated” statistical maps with comparable spatial similarity level (mean ranging from 0.05 to 0.07, standard deviation ranging from 0.25 to 0.29) was also used to calculate the transcriptional overlap with four inhibitory control tasks.

We used a range of different numbers of genes (*x* from 1 to 2000, with step size 10) as thresholds to identify the top *x* most similar genes. We calculated three kinds of overlap under different thresholds: the first one is “within control” overlap, which is the overlap in gene expression association between two inhibitory tasks (e.g., think/no-think and emotion regulation, or stop-signal and go/no-go); the second one is “inhibitory & DMN” overlap, which is the transcriptional overlap between one of the control-related tasks and DMN (e.g., think/no-think and DMN, or emotion regulation and DMN). The third one is the “inhibition & permutation” overlap, which is the overlap of associated gene lists between one of the actual statistical maps (i.e., emotion regulation or memory control) and one of the permutated statistical maps. Finally, we averaged the percentage of each kind of overlap for a certain threshold x and compared the average percentages across all thresholds.

### Gene Ontology Enrichment Analysis

The Gene Ontology(GO) is a widely used bioinformatics tool to interpret the complex gene list based on the knowledge regarding functions of genes and gene products (Ashburner *et al.*, 2000; Huang *et al.*, 2009). To systematically investigate the biological meaning of the “inhibitory- related” genes, we use GO to perform a binary version of the overrepresentation test for Biological Processes (BP), Molecular Function (MF), and Cellular Component (CC). We did not use additional parameters to restrict the selection of Go categories. Currently, experimental findings from over 140 000 published papers are represented as over 600 000 experimentally-supported GO annotations in the GO knowledgebase. For the input gene list, “Gene Ontology enrichment analysis” can identify relevant groups of genes that function together, and associate and statistically test the relationship between these groups and Go annotations. In this way, it can reduce a big list of genes to a much smaller number of biological functions (BP, MF, or CC), and make the input list more easily comprehendible. This method has been applied successfully to understand the output of many gene expression studies, including the study that also used the Allen Human Brain Atlas (Richiardi *et al.*, 2015). More specifically, we used GOATOOLS (Klopfenstein *et al.*, 2018) to perform the GO analyses based on Ontologies and Associations downloaded on 15^th^ November 2018. All of the significant GO items were corrected by FDR correction (p<0.05). We further identified the frequently seen words within all the significant GO items by counting the frequency of the words after removal of not meaningful words (e.g. “of” and “the”).

### Disease-related gene set enrichment

Although GO analysis can provide insights into the biological functions (BP, MF, or CC) of the overlapped gene list, the approach does not provide sufficient information to identify disease associations of the gene list. We leveraged *ToppGene* (Chen *et al.*, 2009) to explore the gene-diseases associations. The ToppGene platform can cluster groups of genes together according to their disease associations, and perform a statistical test as well as the multiple testing error corrections. The latest version of DisGeNET included associations between 17,549 genes and 24,166 diseases, disorders, or abnormal human phenotypes. In this study, our analysis was based on one of the sub-database of DisGeNET, the DisGeNET Curated (expert-curated associations obtained from UniProt and CTD human datasets). We did not use additional parameters to restrict the selection of disease items and performed a binary version of the over-representation analysis. All of the significant diseases items were FDR-correction (p<0.05).

### Data and code availability

All the data (excluding neuroimaging data) are stored in the Open Science Framework (OSF). Open access data includes study summary, extracted coordinates for ALE, significantly associated genes for each task paradigm, and overlapped gene list(s) (OSF link: https://doi.org/10.17605/OSF.IO/6WZ2J). Neurovault was used to store all the neuroimaging data (e.g., results of ALE from Figure2, MACM from Figure3, and “Top Genes” expression maps from Figure 4) and provide 3D visualization of all the statistical maps. (Neurovault link: https://neurovault.org/collections/4845/). Functional-defined brain parcellation can be found in (https://github.com/ThomasYeoLab/CBIG/tree/master/stable_projects/brain_parcellation/Schaefer2018_LocalGlobal) Other research data supporting reported findings are available from the authors upon request.

**Figure 2.**
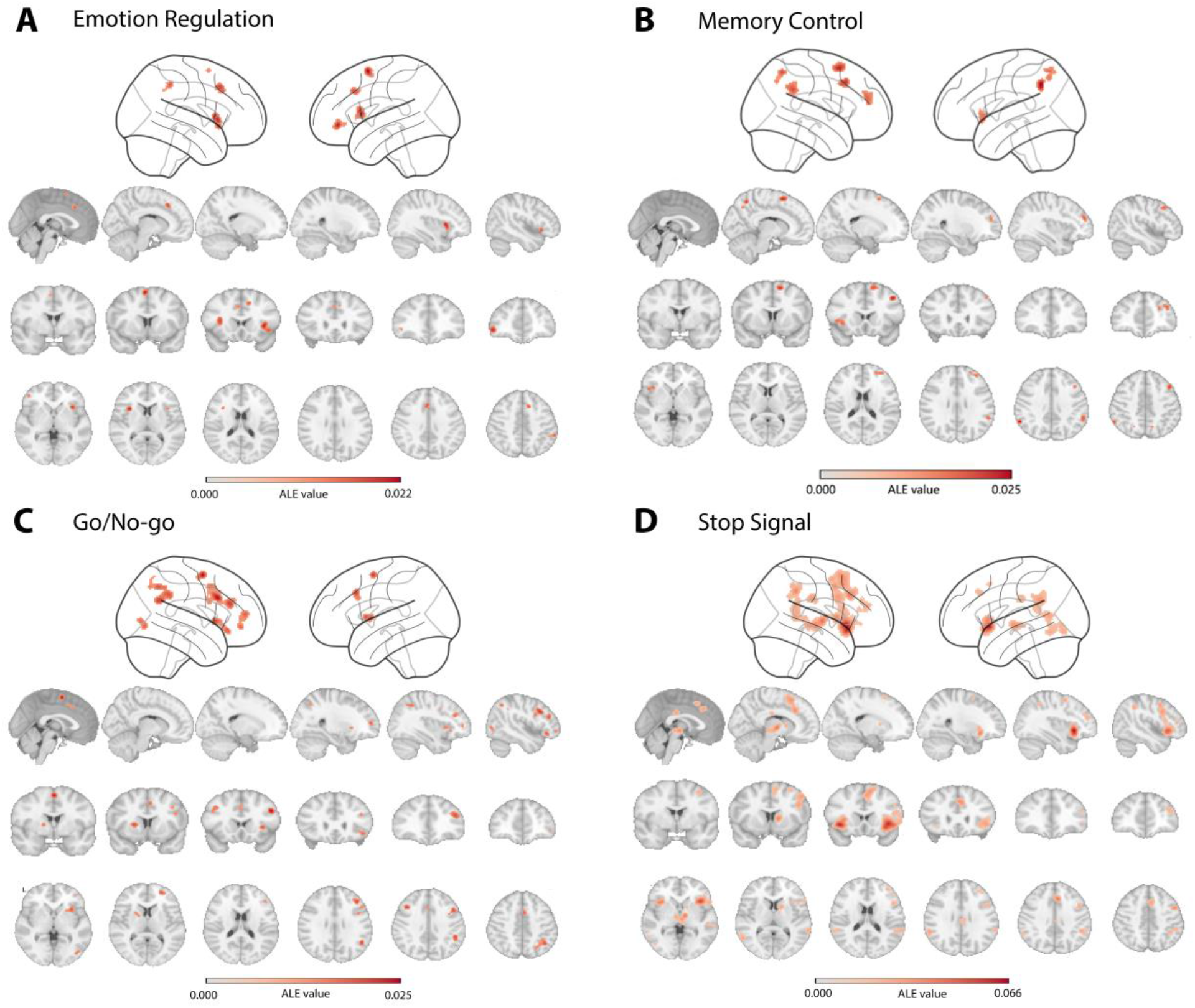
Brain activity underlying four inhibitory control tasks revealed by Activation likelihood estimation (ALE) meta-analyses. (A) Brain regions significantly more activated in the “Regulation” compared to the “Baseline” condition during the Emotion Regulation task. (B) Brain regions significantly more activated in the “Control” compared to the “non-control” condition during the memory control task. (C) Brain regions significantly more activated in the “No-Go” compared to the “Go” condition during the Go/No-go task. (D) Brain regions significantly more activated in the “Stop” compared to the “Go” condition during the Stop-signal task.

**Figure 3.**
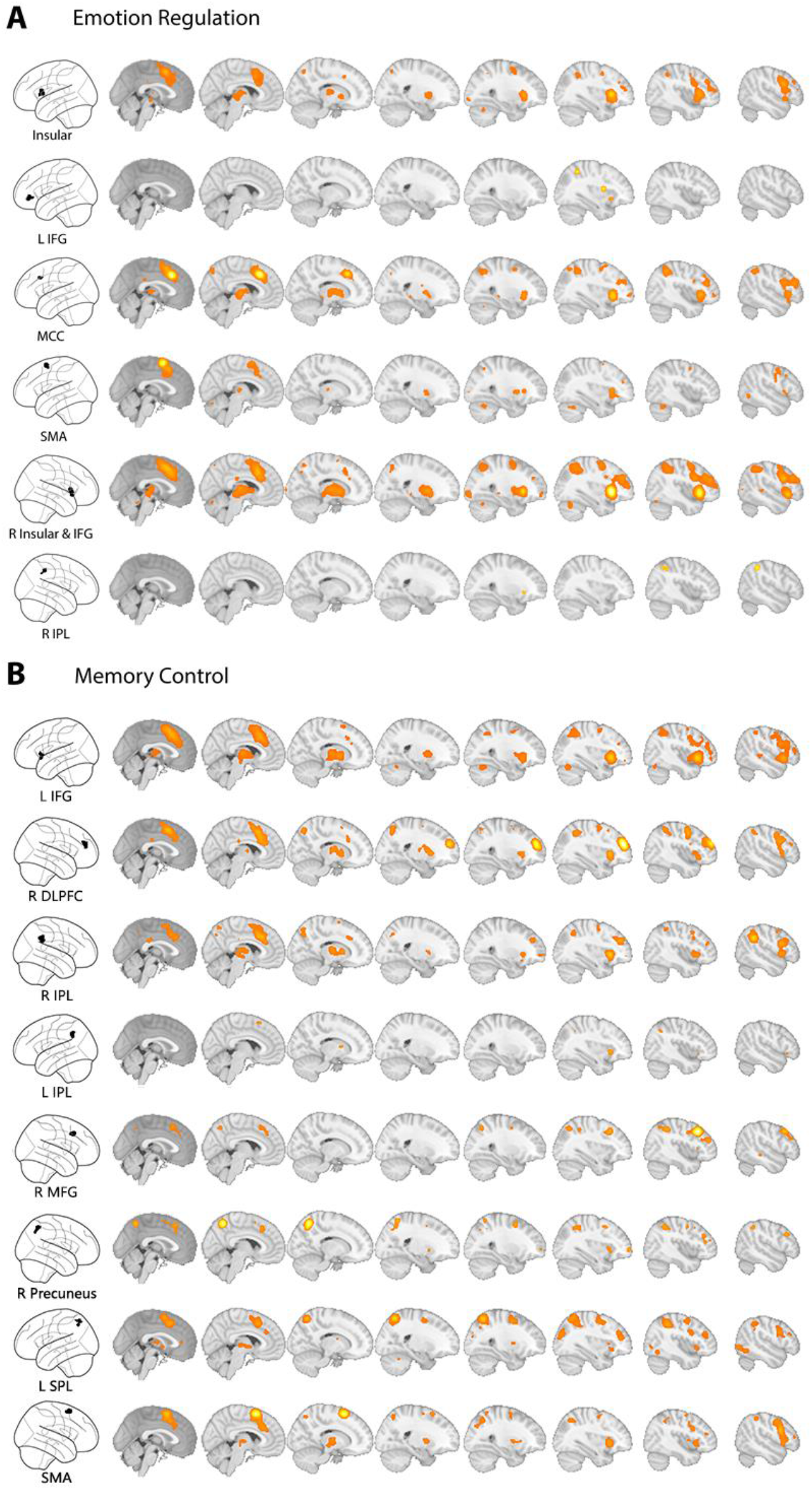
Co-activation maps of Emotion Regulation and Memory Control revealed by Meta-analytic connectivity modeling (MACM). (A) Co-activation maps for regions of interest (ROIs) resulting from ALE analysis of the emotion regulation. (B) Co-activation maps for ROIs resulting from ALE analysis of the memory control. ALE-based ROIs are projected onto glass brains (left column), and co-activation patterns are rendered on an MNI template second to ninth column (right columns)

**Figure 4.**
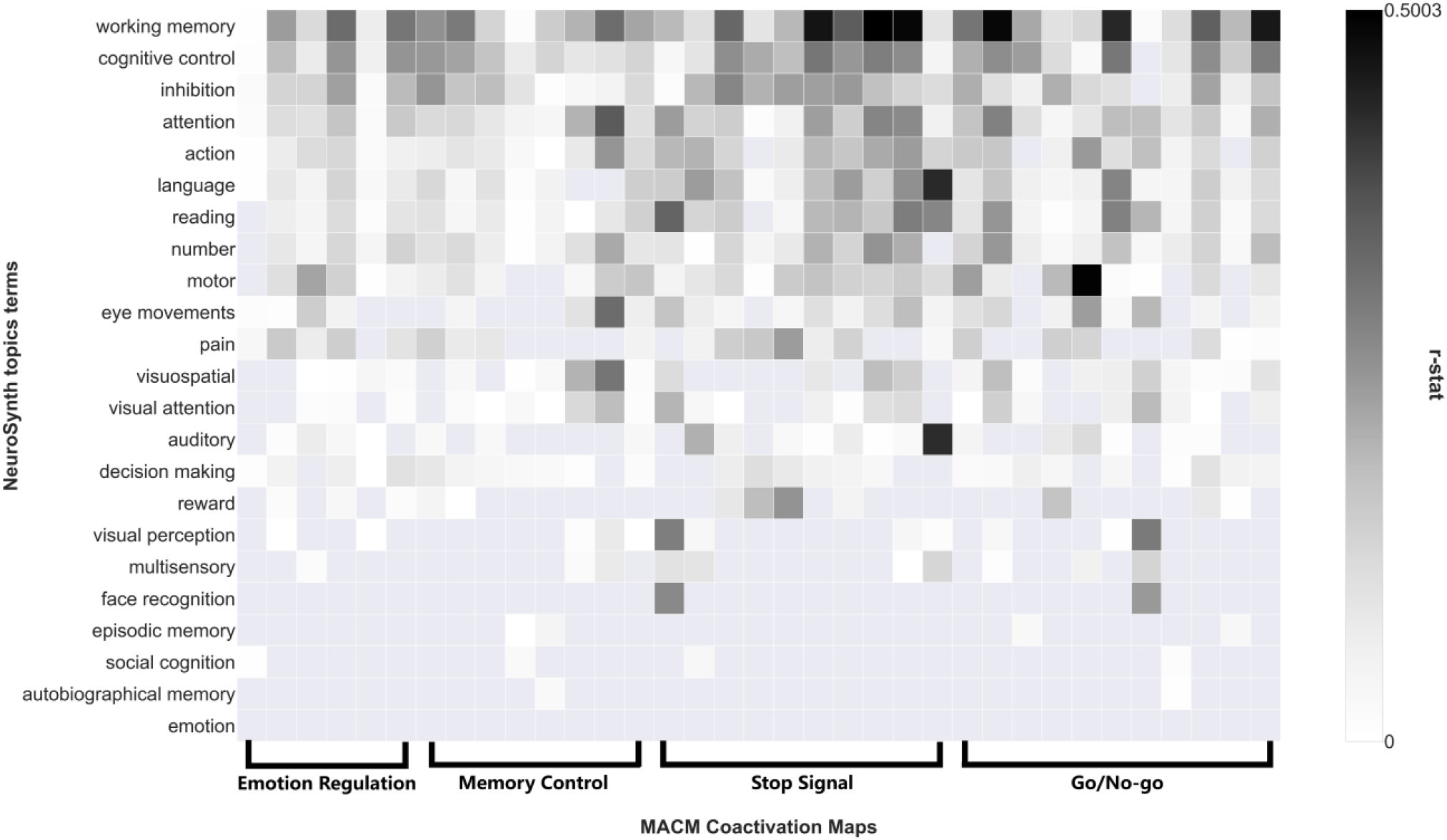
Behavioral profiles of co-activation maps across four inhibitory control tasks. Spatial similarity between Neurosynth meta-maps and co-activation maps across 23 topic terms. Terms are ordered by the mean r-values across the row(s) (all co-activation maps). “Working memory”, “cognitive control”, and “inhibition” are located at the top, suggesting the common stronger association. Domain-specific cognitive functions (e.g. autobiographical memory, emotion) are located at the bottom, suggesting a limited association.

For the neuroimaging meta-analysis, two software (GingerALE and Sleuth) (http://www.brainmap.org/software.html) were used. The Neurosynth Tool (https://github.com/neurosynth/neurosynth), a Python package that covered most of the functions on the Neurosynth (http://www.neurosynth.org/) website, was used in functional profile analyses and co-activation analyses (*see Supplementary Materials*). Activation-gene expression association analyses were based on alleninfo tool (https://github.com/chrisfilo/alleninf) and web application of gene decoder within the Neurovault (https://neurovault.org/). Nilearn (https://nilearn.github.io/) was used to load, manipulate, and visualize MRI statistical maps.

GOATOOLS (https://github.com/tanghaibao/goatools), a Python library, was used for Gene Ontology analyses and related visualization. The Ontology data was downloaded from the Gene Ontology website (http://geneontology.org/ontology/), and the Association data was downloaded from the National Center for Biotechnology Information (ftp://ftp.ncbi.nlm.nih.gov/gene/DATA/). ToppGene Suite (https://toppgene.cchmc.org/) was used for disease-related gene set enrichment. The gene-disease association database can be downloaded from http://www.disgenet.org/.

Anaconda (https://www.anaconda.com/) Python 3.6 version for Win10 was used as the platform for all the programming and statistical analyses. Custom python scripts were written to perform all analyses described based on the mentioned Python packages and be released via the OSF.

## Results

### Regional brain activity associated with emotion regulation and memory control

Taking data from 95 published studies including in total 1995 subjects, we used 15 Emotion Regulation studies to represent emotion regulation, 15 Think/No-think studies to represent memory control. It is noteworthy that we explicitly only focus on emotion regulation studies using the “suppression” or “distraction” as the regulation techniques rather than the more common “cognitive reappraisal,” because of the conceptual relation with memory control, and equal statistical power between contrasted conditions. Furthermore, 27 Go/No-go studies and 38 Stop Signal studies were used to represent response inhibition. (A list of studies and coordinates are available via the Open Science Framework; Search and inclusion criteria in *Materials and Methods*).

#### Regional brain activity of emotion regulation

The meta-analysis of the emotion regulation studies revealed six brain regions that are active during “regulation” compared to a “passive viewing” condition (FWE-cluster level corrected p<.05, uncorrected p<.001, threshold permutations=1000). Emotion regulation task consistently led to activation in right insula/inferior frontal gyrus (IFG), left IFG, left insula, middle cingulate gyrus, right inferior parietal lobule (IPL) and left supplementary motor area (SMA) (Table 1, Figure 2A).

**Table1.** Significant activated clusters during emotion regulation and think/no-think task

| Task | Brain region | Hemisphere | MNI<br>coordinates | Clust<br>er size | Peak voxel<br>Value |
| --- | --- | --- | --- | --- | --- |
| **Emotion**<br>**Regulation** | Insula/IFG | R | 36 16 6 | 209 | 0.02056 |
|  | IFG | L | -46 44 -4 | 138 | 0.02051 |
|  | Insula | L | -34 16 12 | 156 | 0.02089 |
|  | Middle Cingulate Gyrus | L/R | 10 22 44 | 215 | 0.0189 |
|  | Inferior parietal lobule | R | 60 -42 46 | 90 | 0.01688 |
|  | SMA | L | -4 6 62 | 110 | 0.0217 |
| **Memory**<br>**Control** | IFG/Insula | L | -36 20 -4 | 115 | 0.017 |
|  | DLPFC | R | 36 48 24 | 221 | 0.018 |
|  | Supramarginal Gyrus/IPL | R | 56 -44 32 | 167 | 0.02 |
|  | Supramarginal Gyrus/IPL | L | -58 -52 38 | 125 | 0.025 |
|  | Middle Frontal Gyrus | R | 42 20 42 | 126 | 0.022 |
|  | Precuneus | R | 8 -56 54 | 140 | 0.021 |
|  | Precuneus/SPL | L | -16 -64 56 | 114 | 0.017 |
|  | SMA | R | 12 12 60 | 190 | 0.023 |

IFG: inferior frontal gyrus; SMA: supplementary motor area; DLPFC: dorsolateral prefrontal cortex; IPL: Inferior parietal lobule; SPL: Superior parietal lobule

#### Regional brain activity of memory control

Memory control studies revealed five brain regions during the “No-Think” condition compared to the “Think” condition (FWE-cluster level corrected p<.05, uncorrected p<.001, threshold permutations=1000): left insula/IFG, right dorsolateral prefrontal cortex (DLPFC), right middle frontal gyrus, bilateral IPL/supramarginal gyrus, bilateral precuneus, and SMA (Table 1, Figure 2B).

#### Regional brain activity of response inhibition

We used the same meta-analytical approach for all Go/No-go and Stop-signal studies and found similar significant clusters during the “control” condition compared to “baseline” condition (No-go vs go; Stop vs Go) in the insula, IFG, middle cingulate gyrus, SMA, IPL, and DLPFC (Figure 2C and Figure 2D, Table S1 and Table S2).

#### Analyses of convergence and divergence

To examine the spatial convergence and divergence between emotion regulation and memory control activations, we performed formal conjunction and contrast analysis (FWE-cluster level corrected p<.05, threshold permutations=1000). The conjunction and contrast analysis did not find any reliable clusters after correction. Informal overlap analysis of two thresholded maps (i.e., emotion regulation and memory control) revealed a shared cluster of right IPL (MNI:60/-42/42; BA40).

### Co-activation maps of regions associated with emotion regulation and memory control

To gain deeper insight into the co-activation profiles of brain regions associated with emotion regulation and memory control, we used a large database of fMRI studies (*BrainMap*: http://www.brainmap.org/) and applied meta-analytic connectivity modeling (MACM). This approach reveals brain regions that are consistently activated together with the seed regions resulting from the ALE meta-analyses. To control for specific modeling methods and sizes of seed regions, we also performed similar co-activation analyses using *Neurosynth* (http://neurosynth.org/) based on peak voxel coordinates of each seed region instead of the entire clusters. The two approaches yielded similar co-activation maps for each ROI. In the main test, we only report results from the *BrainMap* analysis (*Results from the Neurosynth can be found in Figure S1 and S2*)

In total, we analyzed six ROIs from the meta-analysis of emotion regulation studies. MACM analyses for the left IFG and right IPL revealed co-activation patterns in the direct vicinity of the seed regions. Co-activation analyses of other seed regions, including the bilateral insula, SMA, MACC revealed co-activation between these ROIs and with the cerebellum, IFG, IPL, thalamus, and DLPFC (Figure 3A). Similarly, we analyzed eight seed regions from the meta-analysis of memory control studies. ROIs tended to be co-activated with each other. Only the MACM analysis for the left IPL ROI identified modest co-activation patterns (Figure 3B). In addition, we investigated the co-activation profiles of response inhibition tasks. The co-activation patterns of 11 ROIs from Go/No-go meta-analysis and ten ROIs from Stop-signal meta-analysis were estimated using the same method. We found a brain network including bilateral DLPFC, insular, IPL, thalamus, SMA, and MACC across the co-activation patterns of ROIs from response inhibition.

### Behavioral profiles of the co-activation maps

To identify the cognition domains most strongly associated with these co-activation maps, we created the behavioral profiles of each co-activation maps using the *Neurosynth* database.

First, we quantified the behavioral profiles of co-activation maps of emotion regulation and memory control. As expected, these two sets of co-activation maps were both strongly characterized by terms such as “inhibition”, “cognitive control”, and “working memory”. Then, the behavioral profiles of Stop-signal and Go/no-go co-activation maps were estimated in the same way. Finally, we investigated the commonalities of behavioral characteristics of all co-activation maps across four paradigms and revealed that these co-activation maps have the highest correlations to terms including “inhibition”, “cognitive control”, and “working memory”, compared to other items (Figure 4).

### Transcriptional signatures underlying the Emotion Regulation and Memory control

Next, to understand the transcriptional correlates that may be associated with brain activity elicited by each task, we combined the Allen Human Brain Atlas (AHBA), a brain-wide atlas of gene expression of postmortem brains and spatial pattern correlation. More specifically, we aimed to identify the genes for which the spatial transcriptional patterns are similar to the spatial pattern of brain activity during a given task.

#### Common transcriptional correlates of brain activity

We used the activation-gene expression association analysis to identify two lists of genes whose spatial patterns correlate with emotion regulation-related brain activity patterns or memory control-related brain activity patterns separately. The top ten genes with the highest spatial similarities are represented in Figure 5A, and their expression patterns in the brain are depicted in Figure 5B. (*Complete gene lists in the OSF repository*). We found a substantial overlap (Figure 5A) between the two identified gene lists for the emotion regulation and memory control activation networks: there are in total 1212 genes (60.6% of all correlated genes) whose expression pattern correlated with the brain activity pattern of both tasks. We evaluated the significance of the overlap by generating a set of null distributions of the overlapping genes under the restriction of spatial similarity. Specifically, two gene lists with identical sizes (list1=2531; list2=1529) were randomly selected from all the genes (N=20787), and then overlapping genes between the two random lists were identified. This procedure was repeated 5000 times. 1718 out of 5000 pairs of randomly sampled genes demonstrated a similar level of spatial similarity as our gene lists of interests. We found that the amount of overlapping genes between the memory control and the emotion regulation was significantly larger than the number of overlapping genes within these 1718 randomly sampled overlapping genes lists across different thresholds (threshold=2000, real overlapping=1212, random overlapping=193.28 ± 12.3, p<0.001) (Results at different thresholds presented in Table S4). Furthermore, the significance of the overlap was estimated by generating 100 pairs of “permutated” statistical maps. 72 out of 100 pairs of genes that were associated with these “permutated” maps were used for the estimation because they showed a comparable level of spatial similarity across genes. The amount of real overlapping genes was significantly higher than the number of overlapping genes within these 72 pairs of genes across different thresholds (threshold=2000, real overlapping=1212, permutated overlapping=667.3 ± 147.39, p<0.01). (Full results in Table S5).

**Figure 5.**
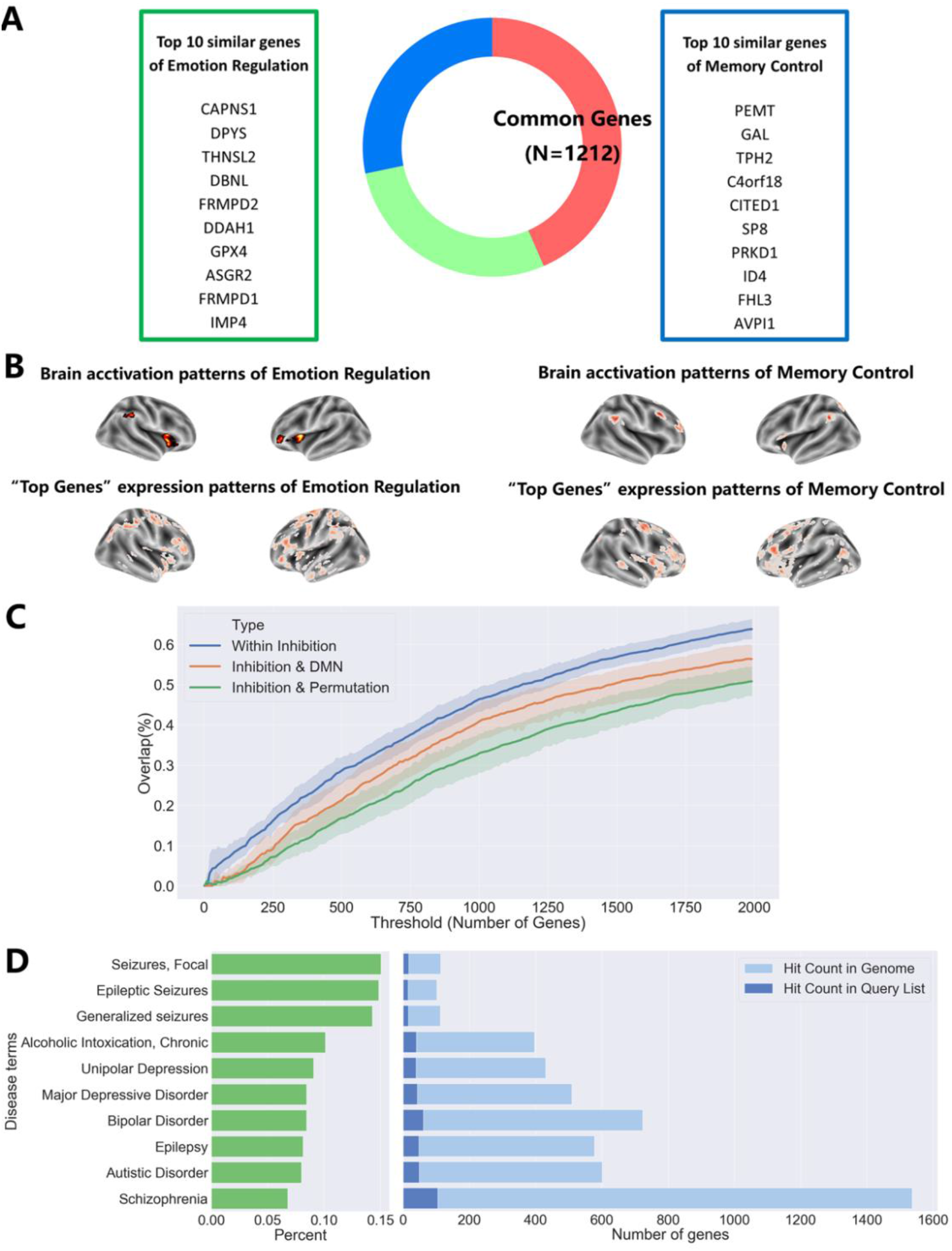
Common transcriptional profiles revealed by activation-gene expression association analysis. (A) Transcriptional patterns of 1212 genes correlated with both the emotion regulation and memory control-related brain activity patterns. (B) Visualization and comparison of brain activation patterns and expression patterns of “Top Genes”. “Top Genes” are 10 genes with the most similar expression patterns as the brain activation. (C) The gene lists of two inhibitory control tasks were more overlap with each other (Within Inhibitory) compared to one of the inhibitory control and DMN (Inhibitory & DMN) or one of the real inhibitory control and one of permutated inhibition inhibitory control (Inhibition & Permutation). (D) Disease associations of “inhibitory-related” genes. “Inhibition-related” genes are associated with genetic risks for depression, bipolar disorder, autism spectrum disorders, schizophrenia, seizures/epilepsy, and chronic alcoholic intoxication. Terms are ordered by the percentage (Left Green) of hit counts in the query list to hit count in the genome. The hit count in the query list is the number of genes belongings to “inhibitory-related” genes (Right Dark Blue). The hit count in the genome is the total number of genes associated with the risk for certain disease terms based on the DisGeNET (Right Light Blue).

#### Specificity of “inhibition-related” genes

To test the specificity of the “inhibition-related” genes versus genes related to neuronal activity and brain function in general, we first generated, also using an ALE meta-analysis, the activation map of “Default Mode Network” (DMN), or “Task-Negative Network” (See Materials and Method; Figure S3). Next, we identified a list of genes associated with the DMN network and “permutated” maps emotion regulation and memory control via the AHBA and the same activation-gene expression association analysis. Given the fact that the DMN has been associated with homeostasis and undirected thought or mind-wandering, we expected that DMN-associated genes are different from inhibition-association genes. To test this, we calculated the number of overlapping genes between all possible combinations of two out of the seven reported gene lists (memory control, emotion regulation, stop-signal, and go/no-go, DMN, “permutated” memory control, “permutated” emotion regulation) across different thresholds. As depicted in Figure 5C, genes associated with inhibitory-control tasks demonstrate more overlap between each other than the associated genes between the inhibitory-control tasks and the DMN (mean Within Inhibition =41.24%, mean Inhibition & DMN =35.18%, t=3.44, p=0.0006). They also showed more overlap compared to associated genes between inhibitory-control tasks and “permutated” inhibitory control maps (mean Within Inhibition =41.24%, mean Inhibition & Permutation =29.53%, t=6.97, p=1.26 ×10-11).

### Biological functions and disease associations of the “inhibition-related” genes

Using gene ontology (GO) (Ashburner *et al.*, 2000), a widely-used literature-based gene-to-function annotation analysis, we generated a list of biological functions related to the “inhibition-related” genes. Furthermore, due to the common finding of impairment in inhibitory control in neurological and psychiatric disorders, we explored the human disease association of the identified gene list.

#### Enrichment for biological functions

Gene Ontology Enrichment Analysis (GOEA) of 779 “inhibition-related” genes identified 5 related Biological Processes (BP), which are associated with information communications between neurons (*Table S6*). More specifically, 19 genes within the list are associated with the neuropeptide signaling pathway, 30 genes with the chemical synaptic transmission, and 17 genes with the potassium ion transmembrane transport, 38 with the cell adhesion, and 60 with signal transduction.

#### Enrichment for human diseases

Additionally, we used *ToppGene* (https://toppgene.cchmc.org/) to perform the gene list enrichment analysis of 708 “inhibition-related” genes for human diseases. Using *DisGeNET* (www.disgenet.org), a comprehensive database on the relationship between human diseases and genes, we found significant associations between our gene list and 36 disease terms (*Table S7*). Among these diseases, seizures/epilepsy, chronic alcoholic intoxication, depression, bipolar disorder, autism spectrum disorders, and schizophrenia were ranked top 10 of the list (i.e., most significant associations) (Figure 5D).

We performed a preliminary investigation of the unique diseases associations for inhibition, but not DMN. First, we calculated the differences between “inhibition-related” genes and DMN-related genes under four different thresholds and included the results into the ToppGene. Only under the threshold of 1500 genes, the algorithm identified a significant association between 103 “inhibition-unique” genes and the risk for Schizophrenia. In other words, variance in these genes, which are associated with inhibition, but not DMN, are reported to be associated with an increased risk for Schizophrenia. We also performed the disease association analysis for “inhibition-related” and DMN-related genes separately. These two sets of genes were associated with similar brain disorders (e.g., Schizophrenia, Autistic Disorder, Unipolar Depression, Alcoholic Intoxication) (See OSF folder). One important difference between the two sets of genes is that “DMN-related” genes are uniquely associated with Alzheimer’s Disease and Age-Related Memory Disorders (Figure S4), which is consistent with our knowledge regarding the relationship between DMN, aging, and memory.

## Discussion

Inhibitory control is a fundamental cognitive function supporting other processes like emotion regulation and response inhibition. Here, we provide neurobiological evidence from human neuroimaging and transcriptional mapping to support the concept that there is one generic neural network of inhibitory control with a set of “inhibition-related” genes modulating not only response inhibition but also emotion regulation and memory control. Our meta-analysis of 95 neuroimaging studies revealed a common role of the right inferior parietal lobule and related regions in a frontal-parietal-insular network during emotion regulation and memory control. These co-activation patterns were also similar to the meta-analysis results of response inhibition tasks, and “inhibition-related” network reported in the literature. Additionally, we used the Allen Human Brain Atlas as an avenue to link this neural network to common transcriptional profiles and identified “inhibition-related” genes, which are associated with the neuronal transmission, and risk for major psychiatric disorders as well as seizures and alcoholic dependency.

The idea that inhibitory control is the underlying core cognitive function in emotion regulation was already suggested before (Ochsner *et al.*, 2002; Ochsner and Gross, 2005; Schmeichel *et al.*, 2008; Wager *et al.*, 2008; McRae *et al.*, 2012). Similarly, it was also suggested that inhibitory control plays a fundamental role in memory control (Levy and Anderson, 2002). A unified theory proposed a central role of inhibitory control in various psychological domains (e.g., motor inhibition, emotional response, and memory retrieval) depending on external task requirements (Aron *et al.*, 2004; Aron *et al.*, 2014; Depue *et al.*, 2016). Engen and Anderson recently reviewed behavioral and neuroimaging studies in this field and proposed the conceptual link between emotion regulation and memory control (Engen and Anderson, 2018). However, there has been little empirical support for this link. Our multimodal analysis provides rich evidence beyond neuroimaging supporting a conceptual link and suggests that inhibitory control, as well as its underlying neural and transcriptional correlates, modulate both emotion regulation and memory control.

Brain activation patterns of emotion regulation and memory control found here are in accordance with previous meta-analyses (emotion regulation: e.g., Buhle et al., 2014; Kohn et al., 2014; memory control: e.g., Guo et al., 2018). We find one overlapping region between emotion regulation and memory control, the right inferior parietal lobule. Due to the central role of inhibitory control in both tasks, only one overlapping brain region seems surprising at first glance. However, meta-analytic connectivity modeling revealed that other regions (including IFG, insula, preSMA/MACC, IPL) that lacked significantly overlapping activations across emotion regulation and memory control, form a tightly integrated network. Our results suggest that although these regions do not overlap strictly, they belong to the same functional network. Our behavioral profile analysis corroborated this interpretation: both co-activation maps of emotion regulation and memory control, as well as stop-signal and go/no-go paradigms, have comparable behavioral profiles and were characterized by terms like “inhibition,” “cognitive control,” and “working memory.”

We have demonstrated the neural commonality of emotion regulation and memory control. Next, we proceeded to investigate if transcriptional profiles overlap with activity patterns in a similar way. Critically, this study adopted an imaging-genetic approach to investigate the common transcriptional signatures across neural networks of emotion regulation and memory control. Activation-gene expression association analysis revealed a largely overlapping gene list whose expression patterns were similar to the activation patterns. Furthermore, we identified a list of “inhibition-related” genes and characterized their biological function and disease associations. “Inhibition-related” genes were primarily associated with the neuropeptide signaling pathway, chemical synaptic transmission, the potassium ion transmembrane transport, and cell adhesion. One common feature of these biological functions is that they are critical for information communications between neurons. Together with our neuroimaging finding of frontal-parietal-insular network, these genes may act as molecular correlates underlying synchronous brain activity during inhibitory control. Inhibition-related” genes were also associated with risks for several psychiatric disorders (e.g., depression, bipolar disorder, schizophrenia, and autism), seizures, and alcoholic dependence.

Recently, neuroimaging (Goodkind *et al.*, 2015; Sha *et al.*, 2018) and genetic studies (Cross-Disorder Group of the Psychiatric Genomics Consortium., 2013; Anttila *et al.*, 2018; Schork *et al.*, 2019) collectively demonstrated the potential common biological roots across psychiatric disorders. However, common phenotypes across disorders are less well understood. Thus, the National Institute of Mental Health’s Research Domain Criteria (NIMH’s RDoC) (https://www.nimh.nih.gov/research/research-funded-by-nimh/rdoc/index.shtml) summarized several domains of phenotypes, where inhibitory control is a central aspect. Here, our results highlighted the critical role of inhibitory control and its biological underpinning across psychiatric disorders. Firstly, dysfunctional inhibitory control (e.g., impaired response inhibition, lack of emotion regulation, and compromised memory control) is evident across different psychiatric disorders (Magee and Zinbarg, 2007; Price and Mohlman, 2007; Tull *et al.*, 2007; Amstadter, 2008; Falconer *et al.*, 2008; Ehring and Quack, 2010; Erk *et al.*, 2010; Joormann and Gotlib, 2010; Lipszyc and Schachar, 2010; Catarino *et al.*, 2015; Yang *et al.*, 2016; Sacchet *et al.*, 2017). Also, inhibitory control deficits can be further linked to some disorder-specific symptoms (e.g., lack of inhibition of negative thoughts (or rumination) in depression, lack of fear control in anxiety, and failure to avoid retrieval of traumatic memories in PTSD). Secondly, our results suggest an overlap between brain regions (frontal-parietal-insular network) involved in inhibitory control and regions whose structural abnormalities were observed consistently in a variety of psychiatric diagnoses (Goodkind *et al.*, 2015). Thirdly, “inhibition-related” genes, which we identified by spatial transcriptional profiles that overlap with activation patterns of inhibitory control tasks, were also associated with the risks for a variety of psychiatric disorders. Furthermore, although brain disorders such as epilepsy and alcoholic dependence may involve different neurobiological underlies compared to psychiatric disorders, impairment of inhibitory control seems to also be a critical behavioral aspect of the epilepsy (Helmstaedter, 2001; Elger *et al.*, 2004) and alcoholic dependence (Lawrence *et al.*, 2009; Courtney *et al.*, 2013; Papachristou *et al.*, 2013).

Our study has two limitations that should be mentioned. First, our neural commonality analyses were based on fMRI studies only. However, overlap in fMRI activation or co-activation patterns lacks temporal information of the underlying cognitive processes. Electroencephalography (EEG) or Magnetoencephalography (MEG) in humans could provide further confirmatory evidence for the idea of a common cognitive process. Recently, Castiglione and colleagues reported that memory control elicited an electrophysiological signature, increased right frontal beta, which was seen in the Stop Signal task (Castiglione *et al.*, 2019). Follow-up electrophysiological studies or even a meta-analysis of them might confirm the idea of a common electrophysiological signature. Second, the current activation-gene expression association analysis is still preliminary (e.g., low sample size and spatial resolution of the post-mortem data), and without the possibility of testing the specific relationship between expression maps and cognitive function. For example, although our preliminary results demonstrated that “inhibition-specific” genes are associated with the risk for Schizophrenia, while DMN-related genes are more closely linked to Alzheimer’s Disease, we also identified considerable transcriptional overlaps between “inhibition-related” genes and DMN-related genes. The latter may suggested that “inhibition-related” genes, as defined in our study may include both “inhibition-specific” genes and other genes supporting general brain function. However, it is challenging to separate them using methods currently available. Nevertheless, the method already showed great potential when helping to understand basic molecular principles of both the structural and functional connectome (*See review by Fornito et al., 2018*) and it identified molecular mechanisms underlying changes in brain structure or function associated with brain disorders (e.g., autism spectrum disorder (Romero-Garcia *et al.*, 2018), Huntington’s disease (McColgan *et al.*, 2018), schizophrenia (Romme *et al.*, 2017), and Alzheimer’s disease (Grothe *et al.*, 2018)). Results from these studies are consistent with the genetics of neuropsychiatric disorders using conventional methods like genome-wide association studies or animal models. Taken together, although preliminary, neuroimaging-gene expression association analysis has demonstrated its potential to bridge brain structure-function associations and to reveal its underlying molecular processes. To detect more specific associations between spatial transcriptional profiles and neuroimaging data, large sets of postmortem gene-expression data with higher spatial resolution need to be collected. Also, more dedicated analytical methods need to be developed and validated (Arnatkevic̆ iūtė *et al.*, 2019) with new methods that may better control for the effects of domain-general genes that are supporting brain function in general and the bias in gene-set enrichment analyses of brain-wide gene expression data, probably induced by the gene-gene co-expression or autocorrelation (Fulcher *et al.*, 2020).

In summary, our multimodal analysis identified a frontal-parietal-insular neural network and a set of genes associated with inhibitory control across emotion regulation, memory control, and response inhibition. The integrative approach established here bridges between cognitive, neural, and molecular correlates of inhibitory control and can be used to study other higher-level cognitive processes. Our findings may deepen our understanding of emotion regulation and memory control in health and pave the way for better emotion regulation and memory control by targeting the core inhibitory-related network or related molecular targets in patients with such deficit at issue.

## Acknowledgments

We thank Prof. Hans van Bokhoven for comments on the early results of this study, Dr. Yuhua Guo for providing brain activation coordinates of unpublished studies, and Dr. Dace Apšvalka for providing valuable feedback on our preprint.

## Author contributions

The experiment was designed by W.L., G.F., and N.K. Data were collected by W.L., N.P., and N.K. Analyses were performed by W.L. and W.L., N.P. wrote the manuscript. W.L, G.F., and N.K. revised and edited the manuscript. All authors provided essential feedback on the manuscript at different stages, contributed substantially to, and approved the final version of the manuscript.

## References

Amstadter, A. (2008). Emotion regulation and anxiety disorders. Journal of anxiety disorders, 22, 211–21

Anderson, M.C., Green, C. (2001). Suppressing unwanted memories by executive control. Nature, 410, 366

Anderson, M.C., Hanslmayr, S. (2014). Neural mechanism of motivated forgetting. Trends in Cognitive Sciences, 18, Issue:, 1–14

Anderson, M.C., Ochsner, K.N., Kuhl, B., et al. (2004). Neural systems underlying the suppression of unwanted memories. Science, 303, 232–35

Anttila, V., Bulik-Sullivan, B., Finucane, H.K., et al. (2018). Analysis of shared heritability in common disorders of the brain. Science, 360, eaap8757

Arnatkevic̆ iūtė, A., Fulcher, B.D., Fornito, A. (2019). A practical guide to linking brain-wide gene expression and neuroimaging data. NeuroImage, 189, 353–67

Aron, A.R., Robbins, T.W., Poldrack, R.A. (2014). Inhibition and the right inferior frontal cortex: one decade on. Trends in cognitive sciences, 18, 177–85

Aron, A.R., Robbins, T.W., Poldrack, R.A. (2004). Inhibition and the right inferior frontal cortex. Trends in cognitive sciences, 8, 170–77

Ashburner, M., Ball, C.A., Blake, J.A., et al. (2000). Gene Ontology: tool for the unification of biology. Nature genetics, 25, 25

Berto, S., Wang, G.-Z., Germi, J., et al. (2017). Human Genomic Signatures of Brain Oscillations During Memory Encoding. Cerebral Cortex, 28, 1733–48

Buhle, J.T., Silvers, J.A., Wager, T.D., et al. (2014). Cognitive reappraisal of emotion: a meta-analysis of human neuroimaging studies. Cerebral cortex, 24, 2981–90

Castiglione, A., Wagner, J., Anderson, M., et al. (2019). Preventing a Thought from Coming to Mind Elicits Increased Right Frontal Beta Just as Stopping Action Does. Cerebral Cortex, 1–13

Catarino, A., Küpper, C.S., Werner-Seidler, A., et al. (2015). Failing to forget: Inhibitory-control deficits compromise memory suppression in posttraumatic stress disorder. Psychological Science, 26, 604–16

Chen, J., Bardes, E.E., Aronow, B.J., et al. (2009). ToppGene Suite for gene list enrichment analysis and candidate gene prioritization. Nucleic acids research, 37, W305--W311

Courtney, K.E., Ghahremani, D.G., Ray, L.A. (2013). Fronto-striatal functional connectivity during response inhibition in alcohol dependence. Addiction biology, 18, 593–604

Cross-Disorder Group of the Psychiatric Genomics Consortium. (2013). Identification of risk loci with shared effects on five major psychiatric disorders: a genome-wide analysis. The Lancet, 381, 1371–79

Depue, B.E., Orr, J.M., Smolker, H.R., et al. (2016). The Organization of Right Prefrontal Networks Reveals Common Mechanisms of Inhibitory Regulation Across Cognitive, Emotional, and Motor Processes. Cerebral Cortex, 26, 1634–46

Dosenbach, N.U.F., Fair, D.A., Miezin, F.M., et al. (2007). Distinct brain networks for adaptive and stable task control in humans. Proceedings of the National Academy of Sciences, 104, 11073–78

Duncan, J. (2010). The multiple-demand (MD) system of the primate brain: mental programs for intelligent behaviour. Trends in cognitive sciences, 14, 172–79

Ehring, T., Quack, D. (2010). Emotion regulation difficulties in trauma survivors: The role of trauma type and PTSD symptom severity. Behavior therapy, 41, 587–98

Eickhoff, S.B., Bzdok, D., Laird, A.R., et al. (2011). Co-activation patterns distinguish cortical modules, their connectivity and functional differentiation. Neuroimage, 57, 938–49

Eickhoff, S.B., Laird, A.R., Grefkes, C., et al. (2009). Coordinate-based activation likelihood estimation meta-analysis of neuroimaging data: A random-effects approach based on empirical estimates of spatial uncertainty. Human brain mapping, 30, 2907–26

Elger, C.E., Helmstaedter, C., Kurthen, M. (2004). Chronic epilepsy and cognition. The Lancet Neurology, 3, 663–72

Elliott, L.T., Sharp, K., Alfaro-Almagro, F., et al. (2018). Genome-wide association studies of brain imaging phenotypes in UK Biobank. Nature, 562, 210

Engen, H.G., Anderson, M.C. (2018). Memory Control: A Fundamental Mechanism of Emotion Regulation. Trends in Cognitive Sciences, 22, 982–95

Erk, S., Mikschl, A., Stier, S., et al. (2010). Acute and sustained effects of cognitive emotion regulation in major depression. Journal of Neuroscience, 30, 15726–34

Falconer, E., Bryant, R., Felmingham, K.L., et al. (2008). The neural networks of inhibitory control in posttraumatic stress disorder. Journal of psychiatry & neuroscience: JPN, 33, 413

Fedorenko, E., Duncan, J., Kanwisher, N. (2013). Broad domain generality in focal regions of frontal and parietal cortex. Proceedings of the National Academy of Sciences, 110, 16616–21

Fornito, A., Arnatkevičiūtė, A., Fulcher, B.D. (2018). Bridging the Gap between Connectome and Transcriptome. Trends in Cognitive Sciences

Fox, P.T., Lancaster, J.L. (2002). Mapping context and content: the BrainMap model. Nature Reviews Neuroscience, 3, 319

Fulcher, B.D., Arnatkeviciute, A., Fornito, A. (2020). Overcoming bias in gene-set enrichment analyses of brain-wide transcriptomic data. bioRxiv

Goodkind, M., Eickhoff, S.B., Oathes, D.J., et al. (2015). Identification of a common neurobiological substrate for mental illness. JAMA psychiatry, 72, 305–15

Gorgolewski, K., Fox, A., Chang, L., et al. (2014). Tight fitting genes: Finding relations between statistical maps and gene expression patterns. Organization for Human Brain Mapping. Hamburg, Germany

Gorgolewski, K.J., Varoquaux, G., Rivera, G., et al. (2015). NeuroVault. org: a web-based repository for collecting and sharing unthresholded statistical maps of the human brain. Frontiers in neuroinformatics, 9, 8

Gross, J.J., Thompson, R.A. (2007). Emotion regulation: Conceptual foundations.

Grothe, M.J., Sepulcre, J., Gonzalez-Escamilla, G., et al. (2018). Molecular properties underlying regional vulnerability to Alzheimer’s disease pathology. Brain, 141, 2755–71

Guo, Y., Schmitz, T.W., Mur, M., et al. (2018). A supramodal role of the basal ganglia in memory and motor inhibition: Meta-analytic evidence. Neuropsychologia, 108, 117–34

Hariri, A.R., Mattay, V.S., Tessitore, A., et al. (2002). Serotonin transporter genetic variation and the response of the human amygdala. Science, 297, 400–403

Hawrylycz, M.J., Lein, E.S., Guillozet-Bongaarts, A.L., et al. (2012). An anatomically comprehensive atlas of the adult human brain transcriptome. Nature, 489, 391–99

Helmstaedter, C. (2001). Behavioral aspects of frontal lobe epilepsy. Epilepsy & Behavior, 2, 384–95

Huang, D.W., Sherman, B.T., Lempicki, R.A. (2009). Systematic and integrative analysis of large gene lists using DAVID bioinformatics resources. Nature Protocols, 4, 44–57

Joormann, J., Gotlib, I.H. (2010). Emotion regulation in depression: relation to cognitive inhibition. Cognition and Emotion, 24, 281–98

Klopfenstein, D. V, Zhang, L., Pedersen, B.S., et al. (2018). GOATOOLS: A Python library for Gene Ontology analyses. Scientific reports, 8, 10872

Kohn, N., Eickhoff, S.B., Scheller, M., et al. (2014). Neural network of cognitive emotion regulation—an ALE meta-analysis and MACM analysis. Neuroimage, 87, 345–55

Kong, X., Song, Y., Zhen, Z., et al. (2016). Genetic Variation in S100B Modulates Neural Processing of Visual Scenes in Han Chinese. Cerebral Cortex, bhv322

Laird, A.R., Eickhoff, S.B., Fox, P.M., et al. (2011). The BrainMap strategy for standardization, sharing, and meta-analysis of neuroimaging data. BMC research notes, 4, 349

Laird, Angela R, Eickhoff, S.B., Kurth, F., et al. (2009). ALE meta-analysis workflows via the brainmap database: progress towards a probabilistic functional brain atlas. Frontiers in neuroinformatics, 3, 23

Laird, A. R., Eickhoff, S.B., Li, K., et al. (2009). Investigating the Functional Heterogeneity of the Default Mode Network Using Coordinate-Based Meta-Analytic Modeling. Journal of Neuroscience, 29, 14496–505

Laird, A.R., Lancaster, J.J., Fox, P.T. (2005). Brainmap. Neuroinformatics, 3, 65–77

Langner, R., Leiberg, S., Hoffstaedter, F., et al. (2018). Towards a human self-regulation system: Common and distinct neural signatures of emotional and behavioural control. Neuroscience & Biobehavioral Reviews

Lawrence, A.J., Luty, J., Bogdan, N.A., et al. (2009). Impulsivity and response inhibition in alcohol dependence and problem gambling. Psychopharmacology, 207, 163–72

Levy, B.J., Anderson, M.C. (2002). Inhibitory processes and the control of memory retrieval. Trends in cognitive sciences, 6, 299–305

Lipszyc, J., Schachar, R. (2010). Inhibitory control and psychopathology: a meta-analysis of studies using the stop signal task. Journal of the International Neuropsychological Society, 16, 1064–76

Magee, J.C., Zinbarg, R.E. (2007). Suppressing and focusing on a negative memory in social anxiety: Effects on unwanted thoughts and mood. Behaviour Research and Therapy, 45, 2836–49

Margulies, D.S., Ghosh, S.S., Goulas, A., et al. (2016). Situating the default-mode network along a principal gradient of macroscale cortical organization. Proceedings of the National Academy of Sciences, 113, 12574–79

McColgan, P., Gregory, S., Seunarine, K.K., et al. (2018). Brain regions showing white matter loss in Huntington’s disease are enriched for synaptic and metabolic genes. Biological psychiatry, 83, 456–65

McRae, K., Jacobs, S.E., Ray, R.D., et al. (2012). Individual differences in reappraisal ability: Links to reappraisal frequency, well-being, and cognitive control. Journal of Research in Personality, 46, 2–7

Morawetz, C., Bode, S., Derntl, B., et al. (2017). The effect of strategies, goals and stimulus material on the neural mechanisms of emotion regulation: A meta-analysis of fMRI studies. Neuroscience & Biobehavioral Reviews, 72, 111–28

Ochsner, K.N., Bunge, S.A., Gross, J.J., et al. (2002). Rethinking feelings: an FMRI study of the cognitive regulation of emotion. Journal of cognitive neuroscience, 14, 1215–29

Ochsner, K.N., Gross, J.J. (2005). The cognitive control of emotion. Trends in cognitive sciences, 9, 242–49

Papachristou, H., Nederkoorn, C., Havermans, R., et al. (2013). Higher levels of trait impulsiveness and a less effective response inhibition are linked to more intense cue-elicited craving for alcohol in alcohol-dependent patients. Psychopharmacology, 228, 641–49

Power, J.D., Cohen, A.L., Nelson, S.M., et al. (2011). Functional network organization of the human brain. Neuron, 72, 665–78

Price, R.B., Mohlman, J. (2007). Inhibitory control and symptom severity in late life generalized anxiety disorder. Behaviour Research and Therapy, 45, 2628–39

Richiardi, J., Altmann, A., Milazzo, A.-C., et al. (2015). Correlated gene expression supports synchronous activity in brain networks. Science, 348, 1241–44

Robbins, T.W. (2016). Neurochemistry of cognition. Oxford Textbook of Cognitive Neurology and Dementia, 91

Romero-Garcia, R., Warrier, V., Bullmore, E.T., et al. (2018). Synaptic and transcriptionally downregulated genes are associated with cortical thickness differences in autism. Molecular psychiatry, 1–12

Romme, I.A.C.C., de Reus, M.A., Ophoff, R.A., et al. (2017). Connectome disconnectivity and cortical gene expression in patients with schizophrenia. Biological psychiatry, 81, 495–502

Sacchet, M.D., Levy, B.J., Hamilton, J.P., et al. (2017). Cognitive and neural consequences of memory suppression in major depressive disorder. Cognitive, Affective, & Behavioral Neuroscience, 17, 77–93

Schaefer, A., Kong, R., Gordon, E.M., et al. (2017). Local-global parcellation of the human cerebral cortex from intrinsic functional connectivity MRI. Cerebral Cortex, 28, 3095–3114

Schmeichel, B.J., Volokhov, R.N., Demaree, H.A. (2008). Working memory capacity and the self-regulation of emotional expression and experience. Journal of personality and social psychology, 95, 1526

Schork, A.J., Won, H., Appadurai, V., et al. (2019). A genome-wide association study of shared risk across psychiatric disorders implicates gene regulation during fetal neurodevelopment. Nature Neuroscience, 240911

Seidlitz, J., Váša, F., Shinn, M., et al. (2018). Morphometric Similarity Networks Detect Microscale Cortical Organization and Predict Inter-Individual Cognitive Variation. Neuron, 97, 231–247.e7

Sha, Z., Wager, T.D., Mechelli, A., et al. (2018). Common Dysfunction of Large-Scale Neurocognitive Networks across Psychiatric Disorders. Biological psychiatry

Shine, J.M., Breakspear, M., Bell, P.T., et al. (2019). Human cognition involves the dynamic integration of neural activity and neuromodulatory systems. Nature Neuroscience

Tull, M.T., Barrett, H.M., McMillan, E.S., et al. (2007). A preliminary investigation of the relationship between emotion regulation difficulties and posttraumatic stress symptoms. Behavior Therapy, 38, 303–13

Wager, T.D., Davidson, M.L., Hughes, B.L., et al. (2008). Prefrontal-subcortical pathways mediating successful emotion regulation. Neuron, 59, 1037–50

Wang, G.Z., Belgard, T.G., Mao, D., et al. (2015). Correspondence between Resting-State Activity and Brain Gene Expression. Neuron, 88, 659–66

Yang, W., Chen, Q., Liu, P., et al. (2016). Abnormal brain activation during directed forgetting of negative memory in depressed patients. Journal of affective disorders, 190, 880–88

Yarkoni, T., Poldrack, R.A., Nichols, T.E., et al. (2011). Large-scale automated synthesis of human functional neuroimaging data. Nature methods, 8, 665

