## Supplementary Material for "Common neural and transcriptional correlates of inhibitory control underlie emotion regulation and memory control"

**This PDF file includes:**

Supplementary Text

Table S1 to S7

Figure S1 to S4

Correspondence:

Wei Liu

Department of Cognitive Neuroscience

Donders Institute for Brain, Cognition and Behaviour

Radboud University Medical Centre

Trigon Building, Kapittelweg 29

6525 EN Nijmegen, The Netherlands

**1. Comparison between BrainMap and Neurosynth-based co-activation analysis**

Co-activation analysis can be performed based on large-scale databases of fMRI studies such as BrainMap or Neurosynth. During the initial data analysis, we took advantages of the relative strengths and weaknesses of the two methods. We augmented the coordinate-based meta-analysis on the Neurosynth with meta-analytic connectivity modelling (MACM) based on the BrainMap to explore co-activation patterns. The idea behind Neurosynth is similar to MACM (which is described in the main text). The Neurosynth algorithm searches for brain regions which co-activate with input coordinates within the same functional contrast, and summarize the results as a co-activation map. Neurosynth has several advantages over BrainMap: (1) Neurosynth is based on automated text mining, therefore includes a higher number of studies and at the same time decreases the potential selection bias of the users. (2) ROI-based MACM is largely dependent on the size (number of voxels) and the shape of the input ROIs. It could be a potential problem because we included far more studies in the ALE analysis of Stop-signal (SS) paradigm and Go/No-go (GN) paradigm compared to the emotion regulation and memory control, leading to on average larger ROIs for stop signal and go/no-go paradigm. On the contrary, Neurosynth-based co-activation analysis does not depend on the ROIs, and it can also base on the MRI coordinates, providing a control for the effect of different size of ROIs. The disadvantage of Neurosynth compared to BrainMap is that automated text mining does not separate different contrasts or experiments within one article. Even though these differences, both methods yielded highly similar co-activation patterns for each ROIs (Figure S1 and Figure S2), suggesting that the effect of methodology and database on the neural network analysis in our study is negligible. We nevertheless chose presented the results of MACM in the main text.

**2. Data sharing of non-imaging data via the Open Science Framework (OSF)**

We used the OSF database (https://osf.io/6wz2j/) as a venue to share non-imaging data generated within this study.

*2.1 Studies and coordinates used in the meta-analyses*

For each task (e.g. ER: emotion regulation; TNT: think/no-think; GN: go/no-go; SS: stop-signal), an excel file with all coordinates used in the meta-analyses is uploaded to the folder (ALE_coordinates_data) within the OSF.

2.2 *Code*

Custom python scripts used in this study can be found.

*2.3 Details of MACM results*

Within the MACM analyses, we estimated in total co-activation patterns of all ROIs from 4 different tasks. Because of the limitation of the number of figures presented in the supplemental materials, we cannot provide all the detailed results for these analyses here. Instead, we used a python package (atlasreader: <https://github.com/miykael/atlasreader>) to generate coordinate tables and region labels from all co-activation images. All results can be found in an OSF folder (folder name: MACM_results_details, filename: MACM_results_altasreader.zip)

*2.4 Lists of genes*

Complete gene lists, together with the statistical results, is uploaded in one folder (complete_gene_list) within the OSF database. All files started with “gene_decoding” are the genes whose expression patterns correlated with the brain activity pattern elicited by each task of interest. All files started with “overlapped_gene_list” are genes whose expression patterns correlated with two (ER and TNT) or more (ER, TNT, GN, and SS) task-related activity patterns simultaneously. Numbers (e.g. 500, 1000) were thresholds used in the analysis to select the genes with most similar spatial patterns. All files started with “difference” are genes whose spatial patterns correlated with inhibition tasks, but not DMN.

*2.5 Tables generated by gene ontology enrichment analyses*

Tables for enrichment analyses of biological functions (GOEA.zip) or diseases items (disease.zip). Results from different thresholds were presented.

**3. Data sharing and 3D visualization of statistical maps via the Neurovault**

We uploaded all the statistical maps generated within this study to our Neurovault database (<https://neurovault.org/collections/4845/>) for data sharing purpose and 3D, interactive visualization. There are in total of 46 maps within the images collection.

- Images names starting with the task names (e.g. ER) and contrast names (e.g. Regulation vs Baseline) are maps resulting from the ALE meta-analysis.
- Images names ending with threshold methods are corrected ALE images (e.g. ALE C05 1K stands for p<.05, uncorrected p<.001, threshold permutations=1000).
- Images names starting with “MACM” are co-activation maps from the MACM analyses.
- No images from the coordinate-based co-activation analysis using Neurosynth were uploaded.

**Table S1 Significant activations resulting from the meta-analysis of Go/No-go paradigm**

| **Brain region** | **Hemisphere** | **MNI coordinates** | **Cluster size** | **Peak voxel Value** |
| --- | --- | --- | --- | --- |
| Insular/IFG | R | 32 16 2 | 212 | 0.020847 |
| Inferior temporal gyrus | R | 48 -72 -4 | 98 | 0.017735 |
| Superior frontal gyrus | R | 24 54 10 | 182 | 0.018687 |
| Putamen | L | -22 8 6 | 127 | 0.019816 |
| DLPFC | R | 40 34 24 | 180 | 0.021783 |
| IFG, opercular part | R | 46 18 32 | 266 | 0.024922 |
| Angular | R | 50 -54 28 | 238 | 0.021796 |
| DLPFC | L | -40 22 38 | 114 | 0.021692 |
| Middle cingulate gyrus | R/L | 4 10 46 | 98 | 0.017317 |
| Inferior Parietal Lobule | R | 40 -56 44 | 200 | 0.019583 |
| SMA | R/L | 0 0 60 | 95 | 0.024901 |

IFG: inferior frontal gyrus; SMA: supplementary motor area; DLPFC: dorsolateral prefrontal cortex

| **Brain region** | **Hemisphere** | **MNI coordinates** | **Cluster size** | **Peak voxel Value** |
| --- | --- | --- | --- | --- |
| Insular/IFG | R | 36 20 -4 | 2050 | 0.066141 |
| Fusiform | L | -40 -60 -12 | 252 | 0.026161 |
| Insular/IFG | L | -38 18 -6 | 690 | 0.062398 |
| Inferior parietal lobule | R | 48 -44 40 | 1011 | 0.038367 |
| Thalamus | R | 10 -10 2 | 812 | 0.045329 |
| Supramarginal gyrus/ Inferior parietal lobule | L | -58 -46 28 | 601 | 0.032832 |
| DLPFC | R | 36 46 20 | 200 | 0.024995 |
| Middle cingulate gyrus | R/L | 2 -24 30 | 181 | 0.031727 |
| SMA | R | 6 24 34 | 861 | 0.042154 |

**Table S2 Significant activations resulting from the meta-analysis of Stop-signal paradigm**

IFG: inferior frontal gyrus; SMA: supplementary motor area; DLPFC: dorsolateral prefrontal cortex

**Table S3** **Estimation of spatial similarity between gene expression maps.**

| **Threshold\Donor** | **ID9861** | **ID10021** | **ID12876** | **ID14380** | **ID15496** | **ID15697** |
| --- | --- | --- | --- | --- | --- | --- |
| **500** | 0.07(0.24) | 0.03(0.26) | 0.08(0.26) | 0.02(0.28) | 0.06(0.29) | 0.07(0.29) |
| **1000** | 0.07(0.24) | 0.03(0.26) | 0.09(0.26) | 0.03(0.28) | 0.07(0.29) | 0.08(0.29) |
| **1500** | 0.07(0.24) | 0.03(0.26) | 0.09(0.26) | 0.03(0.28) | 0.07(0.29) | 0.08(0.29) |
| **2000** | 0.08(0.24) | 0.03(0.26) | 0.10(0.26) | 0.03(0.28) | 0.07(0.29) | 0.08(0.29) |

Mean (standard deviation)

**Table S4 Estimation of the significance of overlap genes under the restriction of spatial similarity.**

| **Threshold** | **Real overlap** | **Significance** | **Random overlap (mean)** | **Random overlap (standard deviation)** |
| --- | --- | --- | --- | --- |
| **500** | 145 | P<0.001 | 12.01 | 3.32 |
| **1000** | 445 | P<0.001 | 48.35 | 6.62 |
| **1500** | 808 | P<0.001 | 108.89 | 9.7 |
| **2000** | 1212 | P<0.001 | 193.28 | 12.3 |

**Table S5** **Estimation of the significance of overlap genes based on permutated statistical maps.**

| **Threshold** | **Real overlap** | **Significance** | **Permutated overlap (mean)** | **Permutated overlap (standard deviation)** |
| --- | --- | --- | --- | --- |
| **500** | 145 | P<0.01 | 48.62 | 25.14 |
| **1000** | 445 | P<0.01 | 190.41 | 70.43 |
| **1500** | 808 | P<0.01 | 405.04 | 113.70 |
| **2000** | 1212 | P<0.01 | 667.30 | 147.39 |

**Table S6 Gene Ontology Enrichment Analysis results of the “inhibition-related” genes**

| **GO** | **NS** | **enrichment** | **name** | **ratio_in_study** | **ratio_in_pop** | **p_uncorrected** | **depth** | **study_count** | **p_fdr_bh** |
| --- | --- | --- | --- | --- | --- | --- | --- | --- | --- |
| GO:0007218 | BP | e | neuropeptide signaling pathway | 19/779 | 101/20913 | 5,3E-09 | 6 | 19 | 3,7E-05 |
| GO:0007268 | BP | e | chemical synaptic transmission | 30/779 | 239/20913 | 6,2E-09 | 7 | 30 | 3,7E-05 |
| GO:0071805 | BP | e | potassium ion transmembrane transport | 17/779 | 114/20913 | 1,1E-06 | 8 | 17 | 0,00434 |
| GO:0007155 | BP | e | cell adhesion | 38/779 | 463/20913 | 5,3E-06 | 2 | 38 | 0,01603 |
| GO:0007165 | BP | e | signal transduction | 60/779 | 898/20913 | 1,2E-05 | 4 | 60 | 0,02976 |

BP: Biological Processes

**Table S7 Disease associations of the "inhibition-related" genes**

| ID | Name | Source | pValue | FDR B&H | FDR B&Y | Bonferroni | Genes from Input | Genes in Annotation |
| --- | --- | --- | --- | --- | --- | --- | --- | --- |
| C0036341 | Schizophrenia | [DisGeNET Curated](https://toppgene.cchmc.org/output.jsp?userdata_id=ea6b4ed6-3fe4-4b24-8530-d57d51b9b8d0) | 1,92E-07 | 5,15E-04 | 4,37E-03 | 5,15E-04 | [104](https://toppgene.cchmc.org/showQueryTerms.jsp?userdata_id=ea6b4ed6-3fe4-4b24-8530-d57d51b9b8d0&feature=dis&row=0) | [1537](https://toppgene.cchmc.org/showTermDetail.jsp?userdata_id=ea6b4ed6-3fe4-4b24-8530-d57d51b9b8d0&category=Disease&id=C0036341) |
| C0005586 | Bipolar Disorder | [DisGeNET Curated](https://toppgene.cchmc.org/output.jsp?userdata_id=ea6b4ed6-3fe4-4b24-8530-d57d51b9b8d0) | 5,85E-07 | 7,86E-04 | 6,66E-03 | 1,57E-03 | [61](https://toppgene.cchmc.org/showQueryTerms.jsp?userdata_id=ea6b4ed6-3fe4-4b24-8530-d57d51b9b8d0&feature=dis&row=1) | [723](https://toppgene.cchmc.org/showTermDetail.jsp?userdata_id=ea6b4ed6-3fe4-4b24-8530-d57d51b9b8d0&category=Disease&id=C0005586) |
| C0001973 | Alcoholic Intoxication, Chronic | [DisGeNET Curated](https://toppgene.cchmc.org/output.jsp?userdata_id=ea6b4ed6-3fe4-4b24-8530-d57d51b9b8d0) | 4,58E-06 | 4,10E-03 | 3,48E-02 | 1,23E-02 | [40](https://toppgene.cchmc.org/showQueryTerms.jsp?userdata_id=ea6b4ed6-3fe4-4b24-8530-d57d51b9b8d0&feature=dis&row=2) | [396](https://toppgene.cchmc.org/showTermDetail.jsp?userdata_id=ea6b4ed6-3fe4-4b24-8530-d57d51b9b8d0&category=Disease&id=C0001973) |
| C0041696 | Unipolar Depression | [DisGeNET Curated](https://toppgene.cchmc.org/output.jsp?userdata_id=ea6b4ed6-3fe4-4b24-8530-d57d51b9b8d0) | 1,33E-04 | 8,94E-02 | 7,58E-01 | 3,58E-01 | [39](https://toppgene.cchmc.org/showQueryTerms.jsp?userdata_id=ea6b4ed6-3fe4-4b24-8530-d57d51b9b8d0&feature=dis&row=3) | [430](https://toppgene.cchmc.org/showTermDetail.jsp?userdata_id=ea6b4ed6-3fe4-4b24-8530-d57d51b9b8d0&category=Disease&id=C0041696) |
| C0014544 | Epilepsy | [DisGeNET Curated](https://toppgene.cchmc.org/output.jsp?userdata_id=ea6b4ed6-3fe4-4b24-8530-d57d51b9b8d0) | 1,85E-04 | 8,95E-02 | 7,59E-01 | 4,98E-01 | [47](https://toppgene.cchmc.org/showQueryTerms.jsp?userdata_id=ea6b4ed6-3fe4-4b24-8530-d57d51b9b8d0&feature=dis&row=4) | [578](https://toppgene.cchmc.org/showTermDetail.jsp?userdata_id=ea6b4ed6-3fe4-4b24-8530-d57d51b9b8d0&category=Disease&id=C0014544) |
| C1269683 | Major Depressive Disorder | [DisGeNET Curated](https://toppgene.cchmc.org/output.jsp?userdata_id=ea6b4ed6-3fe4-4b24-8530-d57d51b9b8d0) | 2,22E-04 | 8,95E-02 | 7,59E-01 | 5,96E-01 | [43](https://toppgene.cchmc.org/showQueryTerms.jsp?userdata_id=ea6b4ed6-3fe4-4b24-8530-d57d51b9b8d0&feature=dis&row=5) | [509](https://toppgene.cchmc.org/showTermDetail.jsp?userdata_id=ea6b4ed6-3fe4-4b24-8530-d57d51b9b8d0&category=Disease&id=C1269683) |
| C0004352 | Autistic Disorder | [DisGeNET Curated](https://toppgene.cchmc.org/output.jsp?userdata_id=ea6b4ed6-3fe4-4b24-8530-d57d51b9b8d0) | 2,33E-04 | 8,95E-02 | 7,59E-01 | 6,27E-01 | [48](https://toppgene.cchmc.org/showQueryTerms.jsp?userdata_id=ea6b4ed6-3fe4-4b24-8530-d57d51b9b8d0&feature=dis&row=6) | [601](https://toppgene.cchmc.org/showTermDetail.jsp?userdata_id=ea6b4ed6-3fe4-4b24-8530-d57d51b9b8d0&category=Disease&id=C0004352) |
| C0751495 | Seizures, Focal | [DisGeNET Curated](https://toppgene.cchmc.org/output.jsp?userdata_id=ea6b4ed6-3fe4-4b24-8530-d57d51b9b8d0) | 6,19E-04 | 2,08E-01 | 1,76E+00 | 1,66E+00 | [17](https://toppgene.cchmc.org/showQueryTerms.jsp?userdata_id=ea6b4ed6-3fe4-4b24-8530-d57d51b9b8d0&feature=dis&row=7) | [113](https://toppgene.cchmc.org/showTermDetail.jsp?userdata_id=ea6b4ed6-3fe4-4b24-8530-d57d51b9b8d0&category=Disease&id=C0751495) |
| C0234533 | Generalized seizures | [DisGeNET Curated](https://toppgene.cchmc.org/output.jsp?userdata_id=ea6b4ed6-3fe4-4b24-8530-d57d51b9b8d0) | 2,67E-03 | 7,96E-01 | 6,75E+00 | 7,16E+00 | [16](https://toppgene.cchmc.org/showQueryTerms.jsp?userdata_id=ea6b4ed6-3fe4-4b24-8530-d57d51b9b8d0&feature=dis&row=8) | [112](https://toppgene.cchmc.org/showTermDetail.jsp?userdata_id=ea6b4ed6-3fe4-4b24-8530-d57d51b9b8d0&category=Disease&id=C0234533) |
| C4317109 | Epileptic Seizures | [DisGeNET Curated](https://toppgene.cchmc.org/output.jsp?userdata_id=ea6b4ed6-3fe4-4b24-8530-d57d51b9b8d0) | 3,33E-03 | 8,95E-01 | 7,58E+00 | 8,95E+00 | [15](https://toppgene.cchmc.org/showQueryTerms.jsp?userdata_id=ea6b4ed6-3fe4-4b24-8530-d57d51b9b8d0&feature=dis&row=9) | [101](https://toppgene.cchmc.org/showTermDetail.jsp?userdata_id=ea6b4ed6-3fe4-4b24-8530-d57d51b9b8d0&category=Disease&id=C4317109) |
| C0004936 | Mental disorders | [DisGeNET Curated](https://toppgene.cchmc.org/output.jsp?userdata_id=ea6b4ed6-3fe4-4b24-8530-d57d51b9b8d0) | 5,58E-03 | 1,16E+00 | 9,84E+00 | 1,50E+01 | [29](https://toppgene.cchmc.org/showQueryTerms.jsp?userdata_id=ea6b4ed6-3fe4-4b24-8530-d57d51b9b8d0&feature=dis&row=10) | [320](https://toppgene.cchmc.org/showTermDetail.jsp?userdata_id=ea6b4ed6-3fe4-4b24-8530-d57d51b9b8d0&category=Disease&id=C0004936) |
| C0525045 | Mood Disorders | [DisGeNET Curated](https://toppgene.cchmc.org/output.jsp?userdata_id=ea6b4ed6-3fe4-4b24-8530-d57d51b9b8d0) | 8,09E-03 | 1,16E+00 | 9,84E+00 | 2,18E+01 | [28](https://toppgene.cchmc.org/showQueryTerms.jsp?userdata_id=ea6b4ed6-3fe4-4b24-8530-d57d51b9b8d0&feature=dis&row=11) | [309](https://toppgene.cchmc.org/showTermDetail.jsp?userdata_id=ea6b4ed6-3fe4-4b24-8530-d57d51b9b8d0&category=Disease&id=C0525045) |
| C0087169 | Withdrawal Symptoms | [DisGeNET Curated](https://toppgene.cchmc.org/output.jsp?userdata_id=ea6b4ed6-3fe4-4b24-8530-d57d51b9b8d0) | 9,48E-03 | 1,16E+00 | 9,84E+00 | 2,55E+01 | [12](https://toppgene.cchmc.org/showQueryTerms.jsp?userdata_id=ea6b4ed6-3fe4-4b24-8530-d57d51b9b8d0&feature=dis&row=12) | [72](https://toppgene.cchmc.org/showTermDetail.jsp?userdata_id=ea6b4ed6-3fe4-4b24-8530-d57d51b9b8d0&category=Disease&id=C0087169) |
| C0027819 | Neuroblastoma | [DisGeNET Curated](https://toppgene.cchmc.org/output.jsp?userdata_id=ea6b4ed6-3fe4-4b24-8530-d57d51b9b8d0) | 1,15E-02 | 1,16E+00 | 9,84E+00 | 3,09E+01 | [94](https://toppgene.cchmc.org/showQueryTerms.jsp?userdata_id=ea6b4ed6-3fe4-4b24-8530-d57d51b9b8d0&feature=dis&row=13) | [1683](https://toppgene.cchmc.org/showTermDetail.jsp?userdata_id=ea6b4ed6-3fe4-4b24-8530-d57d51b9b8d0&category=Disease&id=C0027819) |
| C0422854 | Gustatory seizure | [DisGeNET Curated](https://toppgene.cchmc.org/output.jsp?userdata_id=ea6b4ed6-3fe4-4b24-8530-d57d51b9b8d0) | 1,26E-02 | 1,16E+00 | 9,84E+00 | 3,38E+01 | [14](https://toppgene.cchmc.org/showQueryTerms.jsp?userdata_id=ea6b4ed6-3fe4-4b24-8530-d57d51b9b8d0&feature=dis&row=14) | [99](https://toppgene.cchmc.org/showTermDetail.jsp?userdata_id=ea6b4ed6-3fe4-4b24-8530-d57d51b9b8d0&category=Disease&id=C0422854) |
| C0751123 | Atonic Absence Seizures | [DisGeNET Curated](https://toppgene.cchmc.org/output.jsp?userdata_id=ea6b4ed6-3fe4-4b24-8530-d57d51b9b8d0) | 1,26E-02 | 1,16E+00 | 9,84E+00 | 3,38E+01 | [14](https://toppgene.cchmc.org/showQueryTerms.jsp?userdata_id=ea6b4ed6-3fe4-4b24-8530-d57d51b9b8d0&feature=dis&row=15) | [99](https://toppgene.cchmc.org/showTermDetail.jsp?userdata_id=ea6b4ed6-3fe4-4b24-8530-d57d51b9b8d0&category=Disease&id=C0751123) |
| C0422853 | Olfactory seizure | [DisGeNET Curated](https://toppgene.cchmc.org/output.jsp?userdata_id=ea6b4ed6-3fe4-4b24-8530-d57d51b9b8d0) | 1,26E-02 | 1,16E+00 | 9,84E+00 | 3,38E+01 | [14](https://toppgene.cchmc.org/showQueryTerms.jsp?userdata_id=ea6b4ed6-3fe4-4b24-8530-d57d51b9b8d0&feature=dis&row=16) | [99](https://toppgene.cchmc.org/showTermDetail.jsp?userdata_id=ea6b4ed6-3fe4-4b24-8530-d57d51b9b8d0&category=Disease&id=C0422853) |
| C0270824 | Visual seizure | [DisGeNET Curated](https://toppgene.cchmc.org/output.jsp?userdata_id=ea6b4ed6-3fe4-4b24-8530-d57d51b9b8d0) | 1,26E-02 | 1,16E+00 | 9,84E+00 | 3,38E+01 | [14](https://toppgene.cchmc.org/showQueryTerms.jsp?userdata_id=ea6b4ed6-3fe4-4b24-8530-d57d51b9b8d0&feature=dis&row=17) | [99](https://toppgene.cchmc.org/showTermDetail.jsp?userdata_id=ea6b4ed6-3fe4-4b24-8530-d57d51b9b8d0&category=Disease&id=C0270824) |
| C0751496 | Seizures, Sensory | [DisGeNET Curated](https://toppgene.cchmc.org/output.jsp?userdata_id=ea6b4ed6-3fe4-4b24-8530-d57d51b9b8d0) | 1,26E-02 | 1,16E+00 | 9,84E+00 | 3,38E+01 | [14](https://toppgene.cchmc.org/showQueryTerms.jsp?userdata_id=ea6b4ed6-3fe4-4b24-8530-d57d51b9b8d0&feature=dis&row=18) | [99](https://toppgene.cchmc.org/showTermDetail.jsp?userdata_id=ea6b4ed6-3fe4-4b24-8530-d57d51b9b8d0&category=Disease&id=C0751496) |
| C4505436 | Generalized Absence Seizures | [DisGeNET Curated](https://toppgene.cchmc.org/output.jsp?userdata_id=ea6b4ed6-3fe4-4b24-8530-d57d51b9b8d0) | 1,26E-02 | 1,16E+00 | 9,84E+00 | 3,38E+01 | [14](https://toppgene.cchmc.org/showQueryTerms.jsp?userdata_id=ea6b4ed6-3fe4-4b24-8530-d57d51b9b8d0&feature=dis&row=19) | [99](https://toppgene.cchmc.org/showTermDetail.jsp?userdata_id=ea6b4ed6-3fe4-4b24-8530-d57d51b9b8d0&category=Disease&id=C4505436) |
| C0234535 | Seizures, Clonic | [DisGeNET Curated](https://toppgene.cchmc.org/output.jsp?userdata_id=ea6b4ed6-3fe4-4b24-8530-d57d51b9b8d0) | 1,26E-02 | 1,16E+00 | 9,84E+00 | 3,38E+01 | [14](https://toppgene.cchmc.org/showQueryTerms.jsp?userdata_id=ea6b4ed6-3fe4-4b24-8530-d57d51b9b8d0&feature=dis&row=20) | [99](https://toppgene.cchmc.org/showTermDetail.jsp?userdata_id=ea6b4ed6-3fe4-4b24-8530-d57d51b9b8d0&category=Disease&id=C0234535) |
| C0751110 | Single Seizure | [DisGeNET Curated](https://toppgene.cchmc.org/output.jsp?userdata_id=ea6b4ed6-3fe4-4b24-8530-d57d51b9b8d0) | 1,26E-02 | 1,16E+00 | 9,84E+00 | 3,38E+01 | [14](https://toppgene.cchmc.org/showQueryTerms.jsp?userdata_id=ea6b4ed6-3fe4-4b24-8530-d57d51b9b8d0&feature=dis&row=21) | [99](https://toppgene.cchmc.org/showTermDetail.jsp?userdata_id=ea6b4ed6-3fe4-4b24-8530-d57d51b9b8d0&category=Disease&id=C0751110) |
| C0022333 | Jacksonian Seizure | [DisGeNET Curated](https://toppgene.cchmc.org/output.jsp?userdata_id=ea6b4ed6-3fe4-4b24-8530-d57d51b9b8d0) | 1,26E-02 | 1,16E+00 | 9,84E+00 | 3,38E+01 | [14](https://toppgene.cchmc.org/showQueryTerms.jsp?userdata_id=ea6b4ed6-3fe4-4b24-8530-d57d51b9b8d0&feature=dis&row=22) | [99](https://toppgene.cchmc.org/showTermDetail.jsp?userdata_id=ea6b4ed6-3fe4-4b24-8530-d57d51b9b8d0&category=Disease&id=C0022333) |
| C3495874 | Nonepileptic Seizures | [DisGeNET Curated](https://toppgene.cchmc.org/output.jsp?userdata_id=ea6b4ed6-3fe4-4b24-8530-d57d51b9b8d0) | 1,26E-02 | 1,16E+00 | 9,84E+00 | 3,38E+01 | [14](https://toppgene.cchmc.org/showQueryTerms.jsp?userdata_id=ea6b4ed6-3fe4-4b24-8530-d57d51b9b8d0&feature=dis&row=23) | [99](https://toppgene.cchmc.org/showTermDetail.jsp?userdata_id=ea6b4ed6-3fe4-4b24-8530-d57d51b9b8d0&category=Disease&id=C3495874) |
| C0751056 | Non-epileptic convulsion | [DisGeNET Curated](https://toppgene.cchmc.org/output.jsp?userdata_id=ea6b4ed6-3fe4-4b24-8530-d57d51b9b8d0) | 1,26E-02 | 1,16E+00 | 9,84E+00 | 3,38E+01 | [14](https://toppgene.cchmc.org/showQueryTerms.jsp?userdata_id=ea6b4ed6-3fe4-4b24-8530-d57d51b9b8d0&feature=dis&row=24) | [99](https://toppgene.cchmc.org/showTermDetail.jsp?userdata_id=ea6b4ed6-3fe4-4b24-8530-d57d51b9b8d0&category=Disease&id=C0751056) |
| C0422855 | Vertiginous seizure | [DisGeNET Curated](https://toppgene.cchmc.org/output.jsp?userdata_id=ea6b4ed6-3fe4-4b24-8530-d57d51b9b8d0) | 1,26E-02 | 1,16E+00 | 9,84E+00 | 3,38E+01 | [14](https://toppgene.cchmc.org/showQueryTerms.jsp?userdata_id=ea6b4ed6-3fe4-4b24-8530-d57d51b9b8d0&feature=dis&row=25) | [99](https://toppgene.cchmc.org/showTermDetail.jsp?userdata_id=ea6b4ed6-3fe4-4b24-8530-d57d51b9b8d0&category=Disease&id=C0422855) |
| C4316903 | Absence Seizures | [DisGeNET Curated](https://toppgene.cchmc.org/output.jsp?userdata_id=ea6b4ed6-3fe4-4b24-8530-d57d51b9b8d0) | 1,26E-02 | 1,16E+00 | 9,84E+00 | 3,38E+01 | [14](https://toppgene.cchmc.org/showQueryTerms.jsp?userdata_id=ea6b4ed6-3fe4-4b24-8530-d57d51b9b8d0&feature=dis&row=26) | [99](https://toppgene.cchmc.org/showTermDetail.jsp?userdata_id=ea6b4ed6-3fe4-4b24-8530-d57d51b9b8d0&category=Disease&id=C4316903) |
| C0422850 | Seizures, Somatosensory | [DisGeNET Curated](https://toppgene.cchmc.org/output.jsp?userdata_id=ea6b4ed6-3fe4-4b24-8530-d57d51b9b8d0) | 1,26E-02 | 1,16E+00 | 9,84E+00 | 3,38E+01 | [14](https://toppgene.cchmc.org/showQueryTerms.jsp?userdata_id=ea6b4ed6-3fe4-4b24-8530-d57d51b9b8d0&feature=dis&row=27) | [99](https://toppgene.cchmc.org/showTermDetail.jsp?userdata_id=ea6b4ed6-3fe4-4b24-8530-d57d51b9b8d0&category=Disease&id=C0422850) |
| C0422852 | Seizures, Auditory | [DisGeNET Curated](https://toppgene.cchmc.org/output.jsp?userdata_id=ea6b4ed6-3fe4-4b24-8530-d57d51b9b8d0) | 1,26E-02 | 1,16E+00 | 9,84E+00 | 3,38E+01 | [14](https://toppgene.cchmc.org/showQueryTerms.jsp?userdata_id=ea6b4ed6-3fe4-4b24-8530-d57d51b9b8d0&feature=dis&row=28) | [99](https://toppgene.cchmc.org/showTermDetail.jsp?userdata_id=ea6b4ed6-3fe4-4b24-8530-d57d51b9b8d0&category=Disease&id=C0422852) |
| C1510472 | Drug Dependence | [DisGeNET Curated](https://toppgene.cchmc.org/output.jsp?userdata_id=ea6b4ed6-3fe4-4b24-8530-d57d51b9b8d0) | 1,30E-02 | 1,16E+00 | 9,84E+00 | 3,49E+01 | [19](https://toppgene.cchmc.org/showQueryTerms.jsp?userdata_id=ea6b4ed6-3fe4-4b24-8530-d57d51b9b8d0&feature=dis&row=29) | [170](https://toppgene.cchmc.org/showTermDetail.jsp?userdata_id=ea6b4ed6-3fe4-4b24-8530-d57d51b9b8d0&category=Disease&id=C1510472) |
| C4317123 | Myoclonic Seizures | [DisGeNET Curated](https://toppgene.cchmc.org/output.jsp?userdata_id=ea6b4ed6-3fe4-4b24-8530-d57d51b9b8d0) | 1,41E-02 | 1,18E+00 | 1,00E+01 | 3,80E+01 | [14](https://toppgene.cchmc.org/showQueryTerms.jsp?userdata_id=ea6b4ed6-3fe4-4b24-8530-d57d51b9b8d0&feature=dis&row=30) | [100](https://toppgene.cchmc.org/showTermDetail.jsp?userdata_id=ea6b4ed6-3fe4-4b24-8530-d57d51b9b8d0&category=Disease&id=C4317123) |
| C0270846 | Epileptic drop attack | [DisGeNET Curated](https://toppgene.cchmc.org/output.jsp?userdata_id=ea6b4ed6-3fe4-4b24-8530-d57d51b9b8d0) | 1,41E-02 | 1,18E+00 | 1,00E+01 | 3,80E+01 | [14](https://toppgene.cchmc.org/showQueryTerms.jsp?userdata_id=ea6b4ed6-3fe4-4b24-8530-d57d51b9b8d0&feature=dis&row=31) | [100](https://toppgene.cchmc.org/showTermDetail.jsp?userdata_id=ea6b4ed6-3fe4-4b24-8530-d57d51b9b8d0&category=Disease&id=C0270846) |
| C0011570 | Mental Depression | [DisGeNET Curated](https://toppgene.cchmc.org/output.jsp?userdata_id=ea6b4ed6-3fe4-4b24-8530-d57d51b9b8d0) | 1,45E-02 | 1,18E+00 | 1,00E+01 | 3,91E+01 | [41](https://toppgene.cchmc.org/showQueryTerms.jsp?userdata_id=ea6b4ed6-3fe4-4b24-8530-d57d51b9b8d0&feature=dis&row=32) | [559](https://toppgene.cchmc.org/showTermDetail.jsp?userdata_id=ea6b4ed6-3fe4-4b24-8530-d57d51b9b8d0&category=Disease&id=C0011570) |
| C0086189 | Drug Withdrawal Symptoms | [DisGeNET Curated](https://toppgene.cchmc.org/output.jsp?userdata_id=ea6b4ed6-3fe4-4b24-8530-d57d51b9b8d0) | 1,70E-02 | 1,34E+00 | 1,14E+01 | 4,57E+01 | [10](https://toppgene.cchmc.org/showQueryTerms.jsp?userdata_id=ea6b4ed6-3fe4-4b24-8530-d57d51b9b8d0&feature=dis&row=33) | [53](https://toppgene.cchmc.org/showTermDetail.jsp?userdata_id=ea6b4ed6-3fe4-4b24-8530-d57d51b9b8d0&category=Disease&id=C0086189) |
| C0270844 | Tonic Seizures | [DisGeNET Curated](https://toppgene.cchmc.org/output.jsp?userdata_id=ea6b4ed6-3fe4-4b24-8530-d57d51b9b8d0) | 1,78E-02 | 1,36E+00 | 1,15E+01 | 4,79E+01 | [14](https://toppgene.cchmc.org/showQueryTerms.jsp?userdata_id=ea6b4ed6-3fe4-4b24-8530-d57d51b9b8d0&feature=dis&row=34) | [102](https://toppgene.cchmc.org/showTermDetail.jsp?userdata_id=ea6b4ed6-3fe4-4b24-8530-d57d51b9b8d0&category=Disease&id=C0270844) |
| C0011581 | Depressive disorder | [DisGeNET Curated](https://toppgene.cchmc.org/output.jsp?userdata_id=ea6b4ed6-3fe4-4b24-8530-d57d51b9b8d0) | 1,86E-02 | 1,36E+00 | 1,15E+01 | 4,99E+01 | [43](https://toppgene.cchmc.org/showQueryTerms.jsp?userdata_id=ea6b4ed6-3fe4-4b24-8530-d57d51b9b8d0&feature=dis&row=35) | [604](https://toppgene.cchmc.org/showTermDetail.jsp?userdata_id=ea6b4ed6-3fe4-4b24-8530-d57d51b9b8d0&category=Disease&id=C0011581) |

**Figure S1 Comparisons between BrainMap-based and Neurosynth-based co-activation map for the emotion regulation studies**

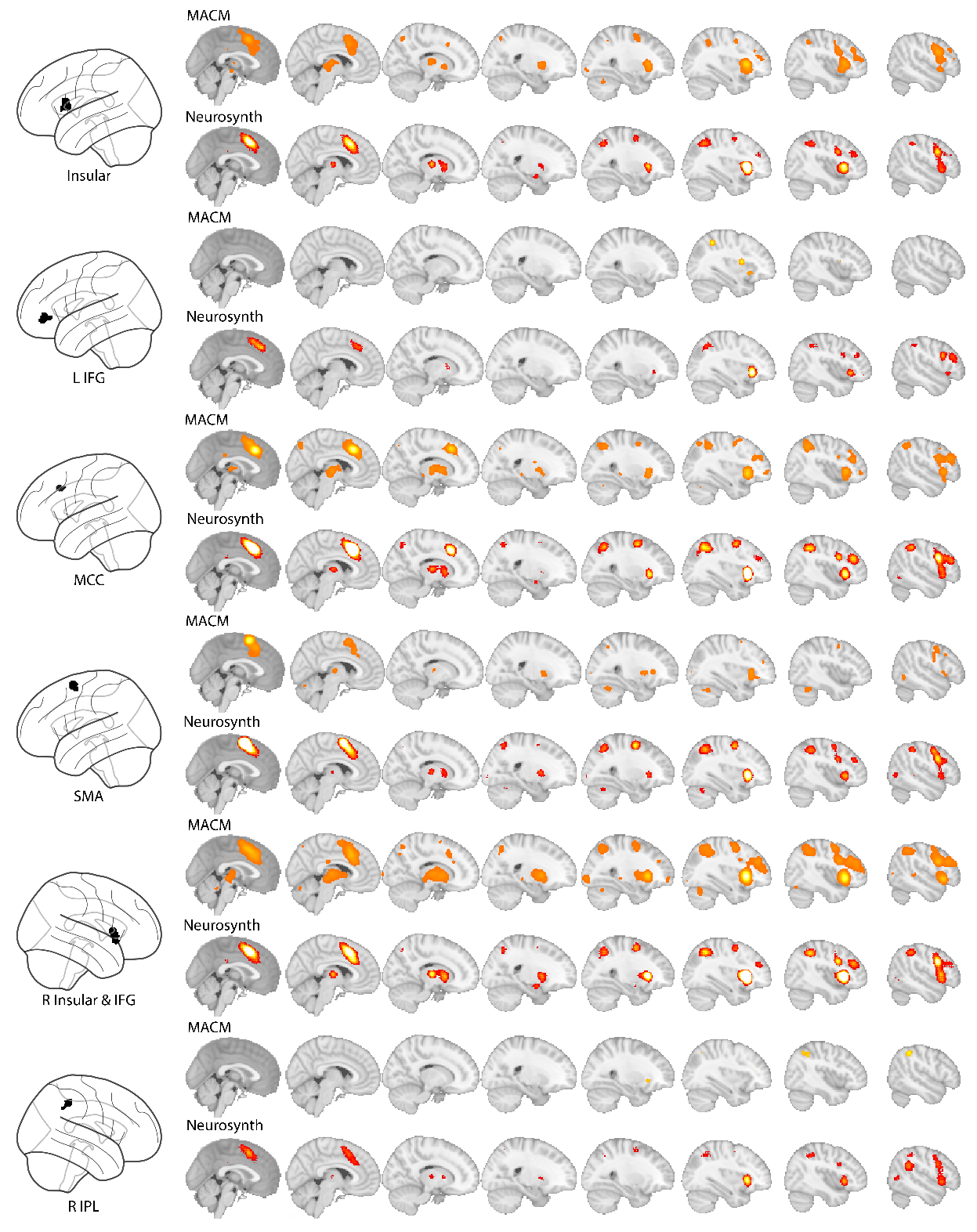

MACM: meta-analytic connectivity modelling; IFG: inferior frontal gyrus; MCC: middle cingulate gyrus; IPL: inferior parietal lobule; SMA: supplementary motor area

**Figure S2 Comparisons between BrainMap-based and Neurosynth-based co-activation map for the memory control studies**

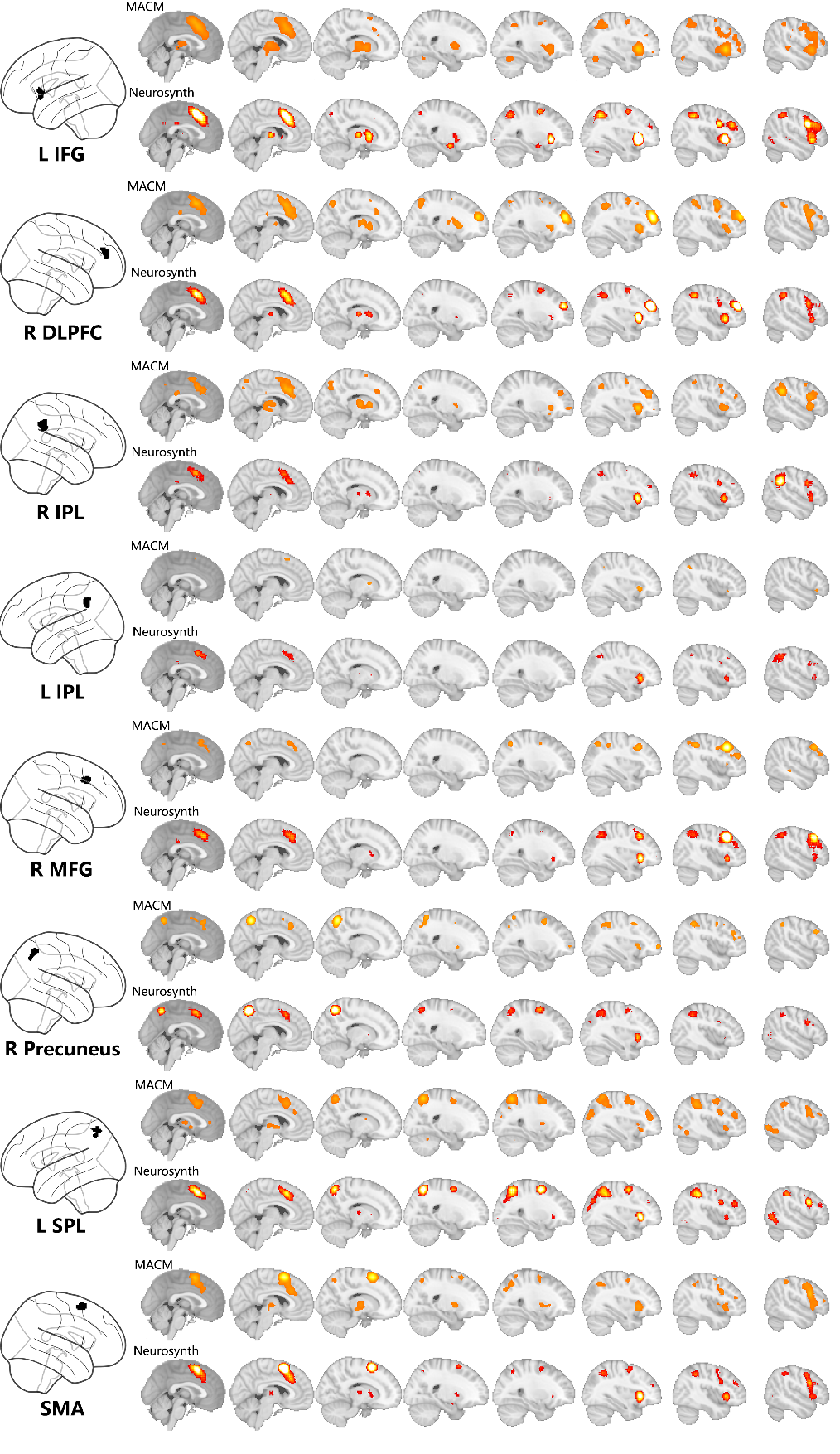

MACM: meta-analytic connectivity modelling; DLPFC: dorsolateral prefrontal cortex; IPL: inferior parietal lobule; SMA: supplementary motor area

**Figure S3 ALE meta-analysis of the Default-Mode Network (Task-negative network)**

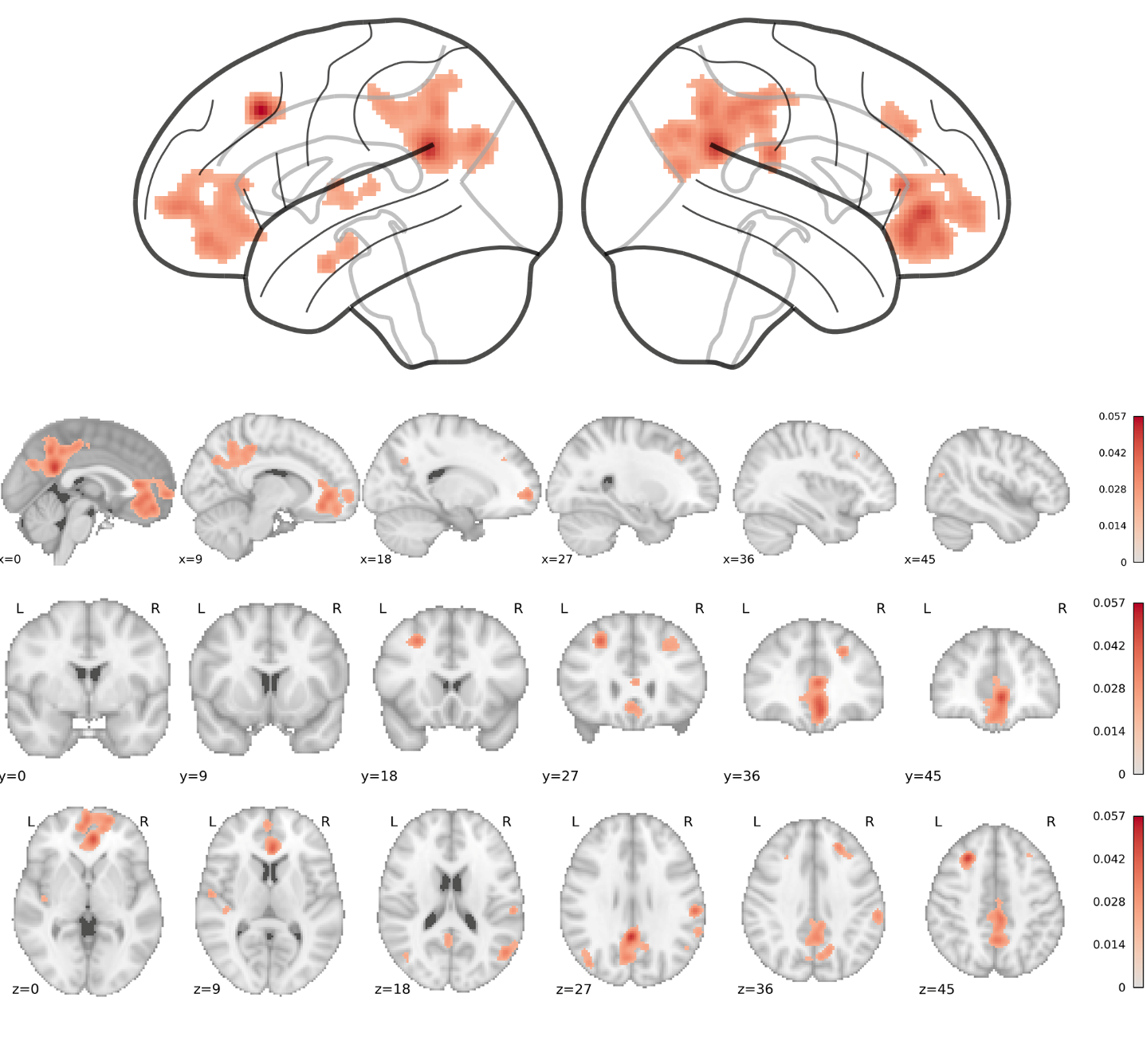

**Figure S4 Comparison of diseases association between “inhibition-related” genes and “DMN-related” genes.**

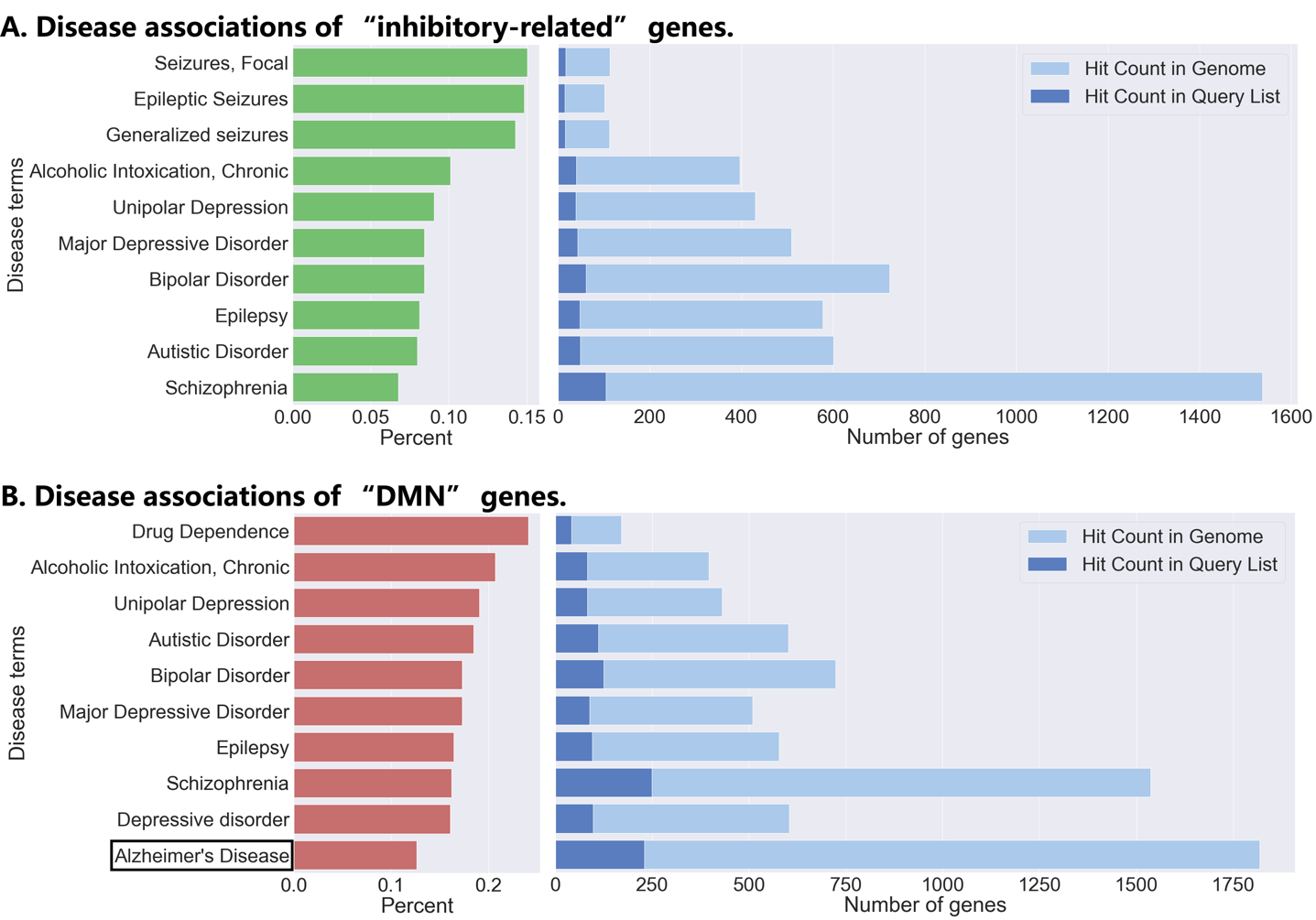
